# Network dynamics underlying activity-timescale differences between cortical regions

**DOI:** 10.1101/2025.09.19.677442

**Authors:** Jose Ernesto Canton-Josh, Lyn A. Ackert-Smith, Renan M. Costa, Lucas Pinto

## Abstract

Network-level dynamics are thought to be central to computation in the cerebral cortex. Yet, how these dynamics differ across areas remains poorly understood. We leveraged an intrinsic property of cortical regions to tackle this problem — the timescales over which they spontaneously sustain activity. We first co-registered functional and spatial transcriptomics datasets to show that timescales across the mouse cortex are predicted by many transcript categories, including those that regulate circuit wiring. Next, we used simultaneous two-photon imaging and optogenetics in mice to ask how these putative differences in connectivity lead to distinct network responses to brief, focal excitatory input to a short-timescale visual area, VISp, and a long-timescale frontal area, MOs. MOs neurons were more likely to respond to photostimulation of their neighbors. Moreover, the evoked dynamics of the overall network were much longer lasting in MOs than VISp, due to the more prevalent recruitment of late-responding neurons, which formed reliable activity sequences. Overall, our findings show that, beyond single-neuron timescales, different cortical areas are distinctly wired to sustain input over varying time windows via network dynamics, with important implications for our understanding of cortical computation.

## Introduction

The mammalian cortex is organized along a functional hierarchy of spontaneous-activity timescales, which are longer in frontal association areas than in posterior sensory regions like the primary visual cortex^1–4^. This organization has attracted increasing attention, given its potential to help explain the differential recruitment of cortical regions, or neurons within them, during cognitive processes like attention, decision making and working memory^5–14^. For instance, the short timescales in sensory regions would be useful for tracking fast-changing features in the environment, whereas longer-lasting activity in association regions could be relevant for accruing or maintaining sensory information in short-term memory over longer time windows^14–17^. Timescale hierarchies are thus emerging as an organizing principle of large-scale cortical dynamics^18^, which makes it crucial to reveal the cellular and circuit mechanisms behind them.

Despite recent efforts, our understanding of these mechanisms is still incomplete. In both primates and rodents, cortical timescale hierarchies covary with multiple cellular and circuit features, e.g. myelination, dendritic density and/or morphology, excitatory and inhibitory neuron subtypes, and expression of glutamate receptor subunits^3,19–23^. However, we do not know the relative contributions of these features in setting regional timescales. In particular, timescales could arise from individual contributions of neurons with long-lasting persistent activity, possibly from intrinsic cellular mechanisms, and/or from more complex interactions between neurons with diverse timescales generating population-level dynamics^8,24–27^. In the latter scenario, more prevalent and/or stronger excitatory synapses in long-than short-timescale regions would lead to stronger reverberatory activity, likely via both local and long-range connections^27–30^. Such areal differences in network dynamics have received much less experimental attention than single-neuron timescales, despite the central role these dynamics play in cortical computation^6,31–37^. In particular, they have not yet been probed with causal perturbation experiments. To address this, we used two-photon optogenetics with near-cellular resolution to compare how two regions of the mouse cortex, primary visual cortex (VISp) and secondary motor cortex (MOs), sustain activity after optical delivery of a brief excitatory input during spontaneous behavior. We found that, while single-neuron response properties were fairly similar between regions, evoked network-level dynamics in MOs outlasted that in VISp due to the recruitment of seconds-long activity sequences. Taken together with a novel analysis of the relationship between cortical timescales and the levels of hundreds of transcripts, our findings suggest that circuit-level mechanisms yield network dynamics on vastly different timescales across the cortex.

## Results

### Cortical timescales are predicted by transcripts regulating circuit wiring

We first asked what transcriptomic features are most predictive of spontaneous timescales across the mouse dorsal cortex on sub-area spatial scales. We reanalyzed our previously collected dataset of cortex-wide mesoscale widefield Ca^2+^ imaging from layer-2/3 excitatory neurons of mice running spontaneously in the dark^14^ (Cux2-Cre x flex-GCaMP6s cross). Briefly, pixel-wise timescales were estimated in the dataset by first removing the contributions of spontaneous running to cortical dynamics using linear regression and then fitting single or dual exponential decay functions to the autocorrelation function of the regression residuals to extract the timescale, *τ*. Importantly, simulations confirmed that this approach accurately recovers the generative timescales in a noisy time series with running-induced modulations in firing rates convolved with Ca^2+^-indicator kernels, evenly and only slightly overestimating timescales across a large range of firing rates and noise levels^14^. Compatible with our previous analysis, we found a gradient of timescales across cortical areas, which was more prominent across the anteroposterior (AP) than the mediolateral (ML) anatomical axes (**Fig. 1A**).

**Figure 1.**
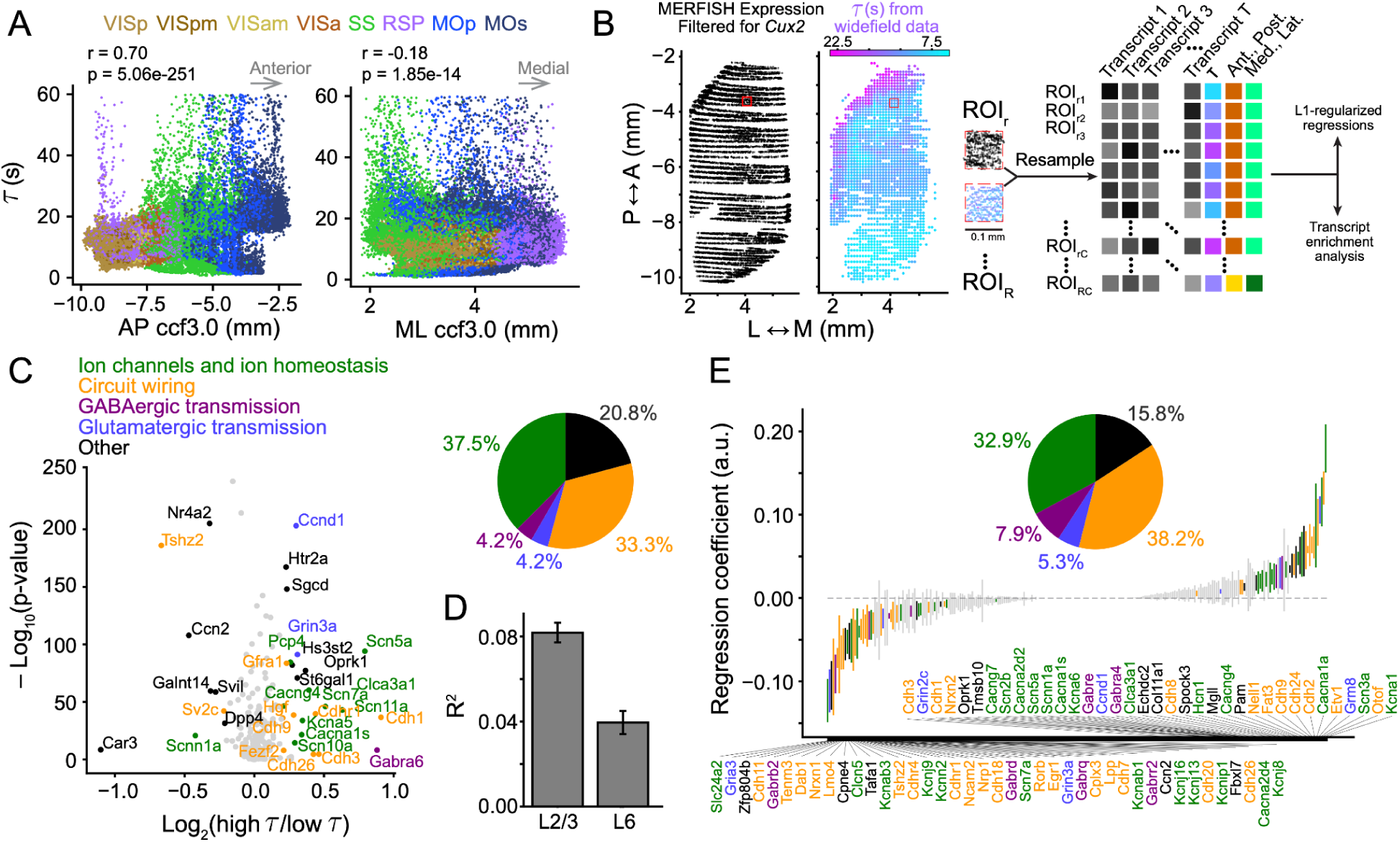
Transcripts implicated in circuit wiring are predictive of timescales in layer-2/3 excitatory neurons. **(A)** Spontaneous timescales as a function of AP and ML position (registered to the Allen ccf3.0), estimated for pixels across dorsal cortical areas (colors). Data were collected using mesoscale widefield Ca^2+^ imaging recordings from mice expressing GCaMP6s in Cux2-expressing excitatory neurons in superficial cortical layers, while they ran spontaneously in the dark (Cux2-Cre x flex-GCaMP6s cross, n = 5 sessions from 5 mice). Printed *r* and p values are from linear correlation analysis. **(B)** Schematics of the analysis used to co-register functional widefield imaging and spatial transcriptomics data and extract transcripts that are enriched in long-timescale pixels or that predict timescales in an L1-regularized regression. **(C)** Volcano plot showing the results of the transcript-enrichment analysis. Non-significant transcripts are shown in gray, significant ones are color-coded according to manually defined categories. Statistical significance was defined as an absolute log_2_(fold-change) larger than 0.2 and a p value < 0.001 (from a two-sided t test, see Methods), with false discovery rate (FDR) correction. Inset: pie chart showing the proportion of significant transcripts in each category. **(D)** Goodness of fit (*R*^2^) for regression models using as predictors transcripts from layer 2/3 (L2/3) Cux2-expressing excitatory neurons and for control models using transcript levels from the same pixels but restricted to layer 6 (L6) excitatory neurons. **(E)** Regression coefficients from the L1-regularized model that predicts pixel-level timescale from transcripts in layer-2/3 *Cux2*-containing neurons. Only significant predictors are labeled, as defined by p < 0.001 with FDR correction (derived from a two-sided t test against zero), and color coded according to the conventions in (C). Inset: pie chart showing the proportion of significant transcripts in each manually defined category. See also Fig. S1, Tables S1 and S2.

To analyze how these timescales are associated with different circuit and cellular features, we used a publicly available spatial transcriptomics dataset from the Allen Institute^38^. We selected a subset of 252 transcripts for ion channels and ion homeostasis-related proteins, neurotransmitter receptor subunits and transporters, synaptic and adhesion proteins, transcription factors and proteins implicated in plasticity and development, and other transcripts used in classifying cortical neuronal subtypes^39^ (**Tables S1, S2**). We then took advantage of two features of our widefield data to align the datasets: dense and broad spatial coverage and laminar specificity. We co-registered the two datasets to the Allen common coordinate framework (ccf3.0), spatially binned them to 100-µm pixels, and filtered the transcriptomics dataset to only include cells with detectable *Cux2* levels, thus matching the genetically defined population in the functional data (**Fig. 1B**). We performed an enrichment analysis to quantify which individual transcripts had significantly more or fewer copy numbers in long-timescale vs. short-timescale pixels across the dorsal cortex (top and bottom quartiles of the *τ* distribution). We found that multiple categories of transcripts were present in higher amounts in long-timescale pixels (**Fig. 1C, Table S1**). For example, compatible with a previous report comparing association and sensory regions^21^, *Grin3a* (glutamate NMDA receptor subunit 3A) is transcribed in higher numbers in these pixels. The same was true for multiple cation channels and a GABA-A receptor subunit, *Gabra6*. Interestingly, many transcripts related to synaptic plasticity and development, such as cadherins^40^, were also present in significantly larger numbers in long-timescales subregions. Overall, transcripts related to ion channel and ion homeostasis comprised 37% of all significant transcripts, while those related to circuit wiring made up 33%.

The analysis above has two potential confounds, however. First, many transcripts are themselves correlated with each other. Second, because timescales covary with anatomical position, the transcripts could be indexing differences in cell-type composition across regions^39^, but not be strictly related to activity timescales. To address this, we devised an L1-regularized regression approach to predict the measured pixel-level *τ* from levels of the 252 transcripts present in single layer-2/3, *Cux2*-containing cells within that pixel (**Fig. 1B**). Moreover, we fitted control models that used transcripts within the same pixels, but taken from layer-6 neurons instead. The layer-2/3 model had higher accuracy, suggesting that it specifically predicts *τ* rather than being trivially related to cortical regions (**Fig. 1D**). We again found that transcripts of many categories are predictive of *τ*, including transcripts for voltage-gated K^+^ and Ca^2+^ channels (*Kcna1, Cacna1*), a metabotropic glutamate receptor (*Grm8*), cadherins (*Cdh2, Cdh24, Ch9*), and transcription factors involved in development (*Etv1, Nell1*) (**Fig. 1E, Table S2**). This analysis revealed that a plurality of transcripts (38%) that predicted timescales are implicated in circuit wiring (**Fig. 1E**), which is compatible with regional differences in dendritic morphology^19,21^. Predicting *τ* after adding ML and AP as model covariates, or selecting transcripts that predicted timescale but not anatomical position yielded compatible results (**Fig. S1**). Thus, beyond previously recognized intrinsic membrane properties and excitatory and inhibitory transmission^21–23^, these results indicate that transcripts involved in circuit wiring may regulate activity timescales. This is consistent with theoretical work showing that stronger recurrent connections in a network can generate longer-lasting dynamics^26,28,29,41^.

### Single-neuron timescales are broadly distributed and spatially clustered in both MOs and VISp, and longer in MOs

The findings above motivated us to investigate how cortical areas potentially differ in terms of their network-level interactions. We first used cellular-resolution two-photon microscopy to compare the dynamics of layer-2/3 excitatory neurons in two areas lying at opposite extremes of the timescale hierarchy — VISp and and the posteromedial portion of MOs^42^ — while mice transgenically expressing GCaMP6s spontaneously ran in the dark (**Fig. 2A–C**, Thy1-GCaMP6s GP4.3 line, n = 1,501 and 2,387 neurons from 4 and 5 mice in VISp and MOs, respectively). We quantified the timescales of single neurons using the same approach used above on widefield data to first regress out running-related activity (**Fig. 2D**). Recapitulating findings from single-unit electrophysiology^2,5,7,12,43,44^, we found broad distributions of timescales in both areas, which were nonetheless overall slightly longer in MOs (**Fig. 2E**, p = 5.5 × 10^-14^, rank-sum test). This was also true when using non-regressed ΔF/F traces (not shown).

**Figure 2.**
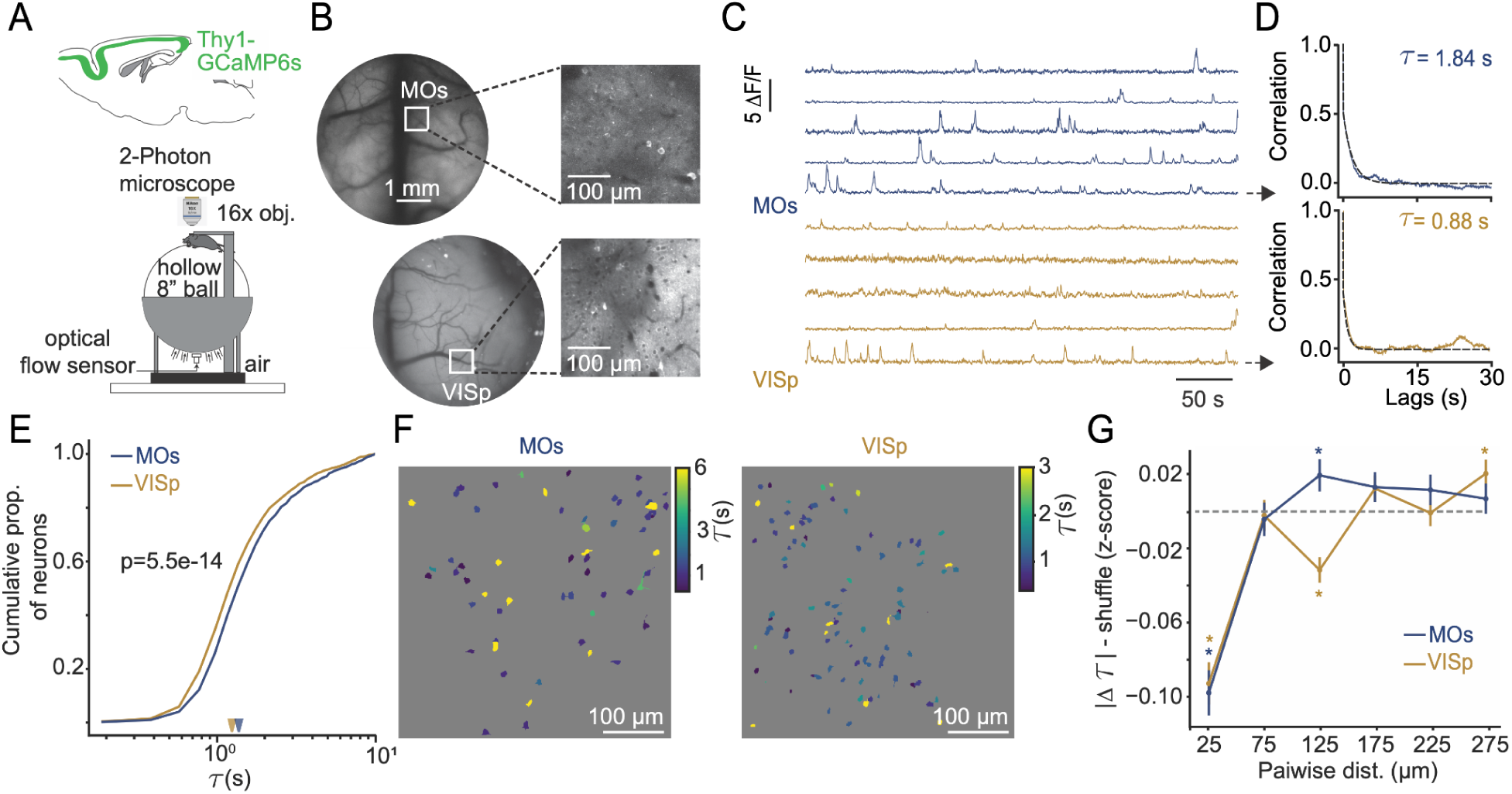
Individual MOs layer-2/3 neurons have longer timescales than those in VISp. **(A)** Schematic of the experimental design. Head-fixed mice expressing the calcium indicator GCaMP6s under the Thy1 promoter were imaged using two-photon microscopy while freely running in the dark on a floating styrofoam ball. **(B)** *Left:* example cranial windows implanted over either the primary visual cortex (VISp) or the secondary motor cortex (MOs), captured with single-photon widefield microcopy. *Right:* example two-photon field of view (FOV) from each region. **(C)** Example ΔF/F traces from individual neurons in MOs (*top*) and VISp (*bottom*). **(D)** Autocorrelation functions and corresponding exponential-decay fits for the example neurons shown by the arrows in (C), used to quantify single-neuron spontaneous timescales *τ*. **(E)** Cumulative distribution of estimated *τ* across all recorded neurons in MOs (n = 1,501 cells from 18 FOVs) and VISp (n = 2,387 cells from 22 FOVs). Arrowheads: medians. P-value is from a rank-sum test. **(F)** Example FOVs from MOs (*left*) and VISp (*right*) with regions of interest (ROIs) color coded according to *τ*. Note spatial clustering of similar colors. **(G)** Shuffle-subtracted absolute pairwise differences in *τ* as a function of their Euclidean distance within the FOV. For both regions, *τ* is more similar than chance for pairs of neurons within ∼75 µm of each other. *: p < 0.05, bootstrap test with FDR correction.

Next, we took advantage of our cellular-resolution imaging approach to investigate whether neurons in each area are spatially organized according to their spontaneous timescale. Initial visual inspection of fields of view suggested that nearby neurons had similar spontaneous timescales, in both MOs and VISp (**Fig. 2F**). To quantify this, for each simultaneously imaged neuronal pair, we computed the difference in timescale and plotted that as a function of physical distance in the tissue, corrected by a shuffling procedure to estimate empirical chance levels of timescale differences (see Methods). Interestingly, we found statistically significant levels of spatial clustering by timescale on the scale of 50–75 µm in both MOs and VISp (**Fig. 2G**, p < 0.05, bootstrap). To our knowledge, this is the first demonstration of spatial clustering by timescale in the cortex. This finding also suggests that timescales are a primary functional property organizing microcircuit layout across the cortex, akin to important coding properties in many cortical regions in the mouse, like orientation selectivity in VISp (e.g., ^45–50^).

### Frontal neurons have more prevalent and widespread responses to brief, focal photostimulation

Autocorrelational measurements of timescales have been proposed to reflect the ability of brain regions/neurons to sustain input over time^2,10^, but this central assumption has not been causally tested. To directly get at this, we used simultaneous two-photon optogenetics and Ca^2+^ imaging^51,52^ (**Fig. 3A**), an approach that has yielded important recent insights into network computation in the cortex^34–37,53^. We did this in the same mice as above, meaning that the spontaneous timescale of every stimulated and responsive region of interest (ROI) was known. Besides constitutive expression of GCaMP6s, we virally co-expressed the red-shifted excitatory opsin ChRmine in cortical excitatory neurons (AAV8-CaMKIIa-ChRmine-mScarlet). While imaging from tens to hundreds of VISp or MOs neurons, we used a separate excitation pathway to briefly optogenetically activate cortical layer 2/3 focally using spiral scanning (15-ms spirals repeated 5 times in 35-ms intervals, total 250 ms), with near-cellular resolution (lateral: ∼20 µm, axial: ∼90–140 µm, **Fig. S2A, B**). Our axial resolution was slightly lower in MOs, presumably due to stronger excitation of the target neuron’s dendritic arbor^54^. However, given that neuronal density is almost double in VISp^55^, we estimate that we directly activated a similar number of neurons in each area (∼6–10). Moreover, optogenetic excitation drove the targeted ROIs in both areas with comparable magnitude and latency (**Fig. 3B–D**, p > 0.1, rank-sum test) and did not depend on the amount of ChRmine expression as quantified from the intensity of the mScarlet fluorescent tag (**Fig. S2C,D**). Thus, we assume that the regional differences reported below are largely due to differences in how this optical input propagates through the circuit downstream of the stimulated neurons. In other words, by directly stimulating cortical sites, we bypass initial stages of thalamocortical or corticocortical transmission to observe how dynamics distinctly evolve after input arrives in layer 2/3 of VISp and MOs (although, as discussed later, we are agnostic as to whether these dynamics are strictly local or additionally involve inter-areal connectivity).

**Figure 3.**
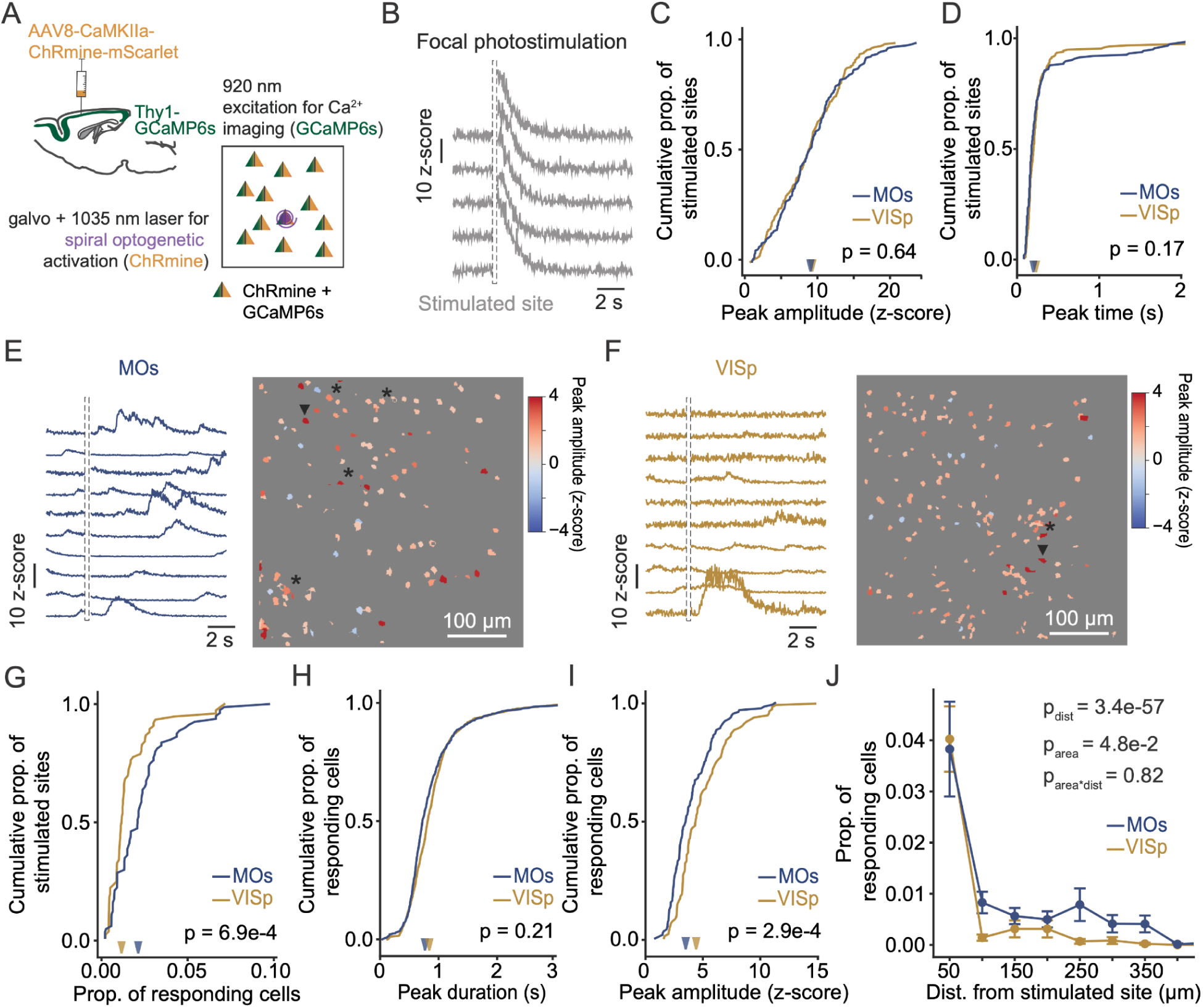
Focal optogenetic stimulation recruits network activity more effectively in MOs than in VISp. **(A)** Schematics of the hybrid transgenic and viral expression strategy and of the simultaneous focal two-photon optogenetic and Ca^2+^ imaging experiment. **(B)** Photostimulation-triggered Ca^2+^ traces from individual trials of a directly stimulated ROI. Dashed outlines correspond to the photostimulation period, when photomultiplier tubes (PMTs) are shuttered. **(C)** Cumulative distribution of peak-response magnitude from directly stimulated sites in VISp (n = 220 from 22 FOVs in 5 mice) and MOs (n = 180 from 18 FOVs in 4 mice). Arrowheads: medians. Printed p-value is from a rank-sum test. **(D)** Same as (C), for time of peak response. **(E)** *Left:* Example single-trial traces for 10 non-stimulated neurons within the same MOs FOV. *Right:* Color-coded peak-response magnitude for all ROIs within the same FOV. Arrowhead: photostimulation site; *: ROIs that showed significant activation. **(F)** Same as (E), but for an FOV in VISp. **(G)** Cumulative distribution of the proportion of significantly responding neurons per photostimulation site. Arrowheads: medians. Printed p-value is from a rank-sum test. **(H)** Same as (G), for peak-response duration (half width at half maximum). **(I)** Same as (G), for peak-response magnitude. **(J)** Relationship between binned distance from the photostimulated site and the probability of a significant neural response for VISp and MOs, showing more spatially widespread responses in MOs. Error bars: ± S.E.M. Printed p-values are from a two-way ANOVA with factors area and distance. See also Fig. S2.

We first compared MOs and VISp in terms of the likelihood that individual neurons became significantly active within 10 s of photostimulation (**Fig. 3E, F**; note that we are interested in network dynamics and do not assume these are monosynaptic interactions). Crucially, for all analyses we excluded ROIs within a 25-µm radius of the photostimulation site, based on our measured lateral resolution. MOs neurons were significantly more likely to become active after focal photostimulation than VISp ones (**Fig. 3G**, p = 6.9 × 10^-4^, rank-sum test, median response probabilities of 0.025 and 0.014 in MOs and V1, respectively). On the other hand, responding MOs cells had peaks of significantly smaller amplitude than VISp ones, with similar duration (**Fig. 3H, I**, p = 2.9 × 10^-4^ and 0.21, respectively, rank-sum test). Moreover, MOs and VISp had similar proportions of neurons that were inhibited by photostimulation (**Fig. S2E**). Finally, the spatial profile of significant responses also differed between areas, with a higher proportion of responsive neurons farther from the photostimulation site in MOs than VISp (**Fig. 3J**). These findings were qualitatively similar for a much briefer photostimulus (**Fig. S2F, G**). Altogether, however, these differences in single-neuron responses to focal photostimulation between MOs and VISp were modest compared to their substantially different area-level timescales (**Fig. 1A**). This prompted us to investigate other potential differences between the two areas.

### Evoked network dynamics are longer lasting in frontal than visual circuits

We next asked whether evoked network-level dynamics differed between MOs and VISp. We first averaged the time course of evoked responses across all significantly responding neurons in each area and observed that average activity remained elevated for longer in MOs than VISp (**Fig. 4A**). To quantify this, we fitted a single exponential decay function to the autocorrelation function of the average evoked responses, computed for many random subsets of neurons, and extracted decay time constants *τ*. Even when we subsampled as few as five neurons, evoked activity in MOs was significantly slower than VISp, which remained consistent as we subsampled more neurons (**Fig. 4B, C**, *τ_MOs_* = 1.72 s ± 0.16 s, *τ_VISp_* = 1.38 s ± 0.01 s, n = 100 neurons per group, mean ± S.D., p < 0.01, bootstrap test). This longer-lasting average network activity in MOs could be due to more sustained responses of individual neurons. However, our analysis of response-peak duration argues against this possibility (**Fig. 3H**). Thus, we also considered the possibility that differences in population dynamics might explain this finding. We first analyzed the distribution of peak times of the evoked responses in the two regions. Peak times occurred significantly later in MOs than VISp (**Fig. 4D, E**, p = 2.8 × 10^-6^, rank-sum test). Indeed, plotting the individual time courses of all significant responses in MOs and VISp revealed that most VISp neurons peaked within 1 s of photostimulation (∼83%). Conversely, these early responding neurons comprised only ∼41% of the MOs population, while most neurons evenly tiled the 10 seconds following photostimulation. These findings were qualitatively similar after the enforcement of additional statistical criteria (**Fig. S3A, B**). Moreover, they could not be ascribed to changes in running behavior, since this was similarly unaffected by phostimulation of each region (**Fig. S3C**). These seconds-delayed responses are highly unlikely to reflect monosynaptic connections to photostimulated neurons. Rather, they are compatible with overall stronger and/or more prevalent connectivity in MOs, as suggested by our analysis of transcriptomics data (**Fig. 1**).

**Figure 4.**
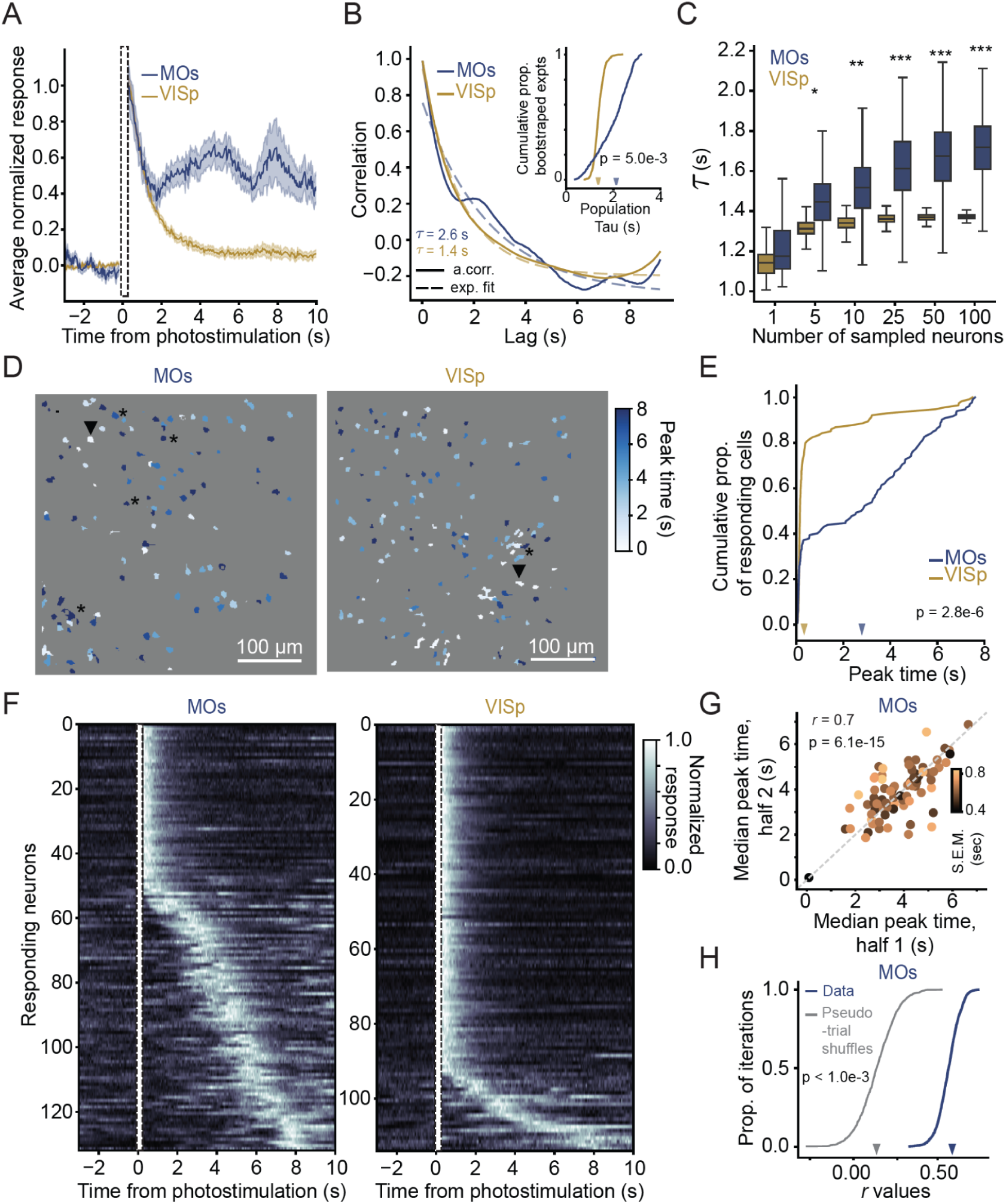
Focal optogenetic stimulation evokes longer-lasting network dynamics in MOs than VISp. **(A)** Peak-normalized photostimulation-triggered responses averaged over all significantly responding neurons in MOs (n = 132 neurons) and VISp (n = 114 neurons). Shaded areas: ± S.E.M. **(B)** Quantification of population-response time constants. Shown are the autocorrelation function and exponential decay fit of the mean response from a random subset of 25 neurons from each cortical area, with extracted *τ* printed. *Inset:* Cumulative distribution of *τ* across 1000 random samples of random 25-neuron subsets. Arrowheads: medians. Printed p-value is from a permutation test. **(C)** Distributions of median *τ* values calculated as in (B) inset, across increasing subset sizes (sampled 1000 times per condition). **(D)** Example FOVs from MOs and VISp with ROIs color-coded according to their peak times.. Arrowheads: stimulated site. *: ROIs that showed significant activation. **(E)** Cumulative distribution of peak times for each area. Arrowheads: medians. Printed p-value is from a rank-sum test. **(F)** Trial-averaged normalized response time courses for all significantly responding neurons in each area, sorted by peak time (note that the same neuron may occur more than once if it responds to more than one photostimulated site). **(G)** Scatter plot showing an example iteration of the cross-validation analysis for peak-time consistency in MOs, where each axis corresponds to the median peak time for non-overlapping halves of a 20-trial set. Data points (neurons) are color coded according to the S.E.M. of peak times over the 20 trials. Printed *r* and *p* are from linear correlation for this iteration. **(H)** Distribution of linear correlation coefficients, *r*, as computed in (G) for 1000 iterations, for the data and pseudo-trials generated from each cell’s spontaneous activity. Arrowheads: medians. Printed p-value is from a permutation test. See also Fig. S3.

The individual response profiles above, particularly in MOs, are reminiscent of activity sequences, which have been previously observed in the context of spontaneous cortical activity^56^ and have been proposed to enable population-level integration and maintenance of information during cognitive tasks^57–65^. We therefore wondered if the average activity patterns above did correspond to sequential activation. To answer this, we performed a subset of experiments in MOs with much higher trial counts, which allowed us to use a cross-validation approach to verify the reliability of peak times across trials. This analysis confirmed that peak times across different sets of trials are highly correlated to each other and significantly higher than expected by chance as assessed by shuffling controls (**Figs. 4F, G;** see **S3D, E** for all experiments in both areas). These findings suggest that frontal region MOs is intrinsically better equipped to generate sequential activity than VISp.

### Evoked dynamics are related to the network’s state at the moment of stimulation

Our findings above suggest that, while photostimulation-evoked activity timing is reproducible (at least in MOs), it also displays trial-to-trial variability. Previous reports have shown that variability in responses to visual stimuli and in motor-related activity in the rodent and primate cortex can be predicted by the initial state of the network, resembling linear dynamical systems^66–68^. To investigate whether similar phenomena explained trial-to-trial variability in our data, we used principal components analysis (PCA) to decompose population activity measured during the initial 5–10-min recording session used to determine spontaneous timescales. We included all non-targeted neurons in the field of view in this analysis, regardless of whether they responded to photostimulation. We then projected the activity of each photostimulation trial onto this space (−3 to 7.5 s from stimulation). Visual inspection of the projections onto the first 3 principal components (PCs) suggested that, when pre-stimulation baselines for different trials were close together in this activity space, their evoked trajectories tended to be more similar than if they started farther apart (**Fig. 5A**). To quantify this, we computed the Euclidean distance between pairs of pre-trial baselines and their respective post-stimulation trajectories on the subspaces spanned by the top 10 PCs (which accounted for 35 ± 2 % and 43 ± 2 % of the variance in VISp and MOs, respectively, mean ± S.E.M., **Fig. 5B**). Pairwise distances between pre-trial baselines were highly correlated with those between post-stimulation trajectories in both areas (**Fig. 5C, D**), and equally so (**Fig. 5E**). Moreover, these correlations were significantly larger than expected by chance as estimated from trial shuffles (**Fig. 5F, G**).

**Figure 5.**
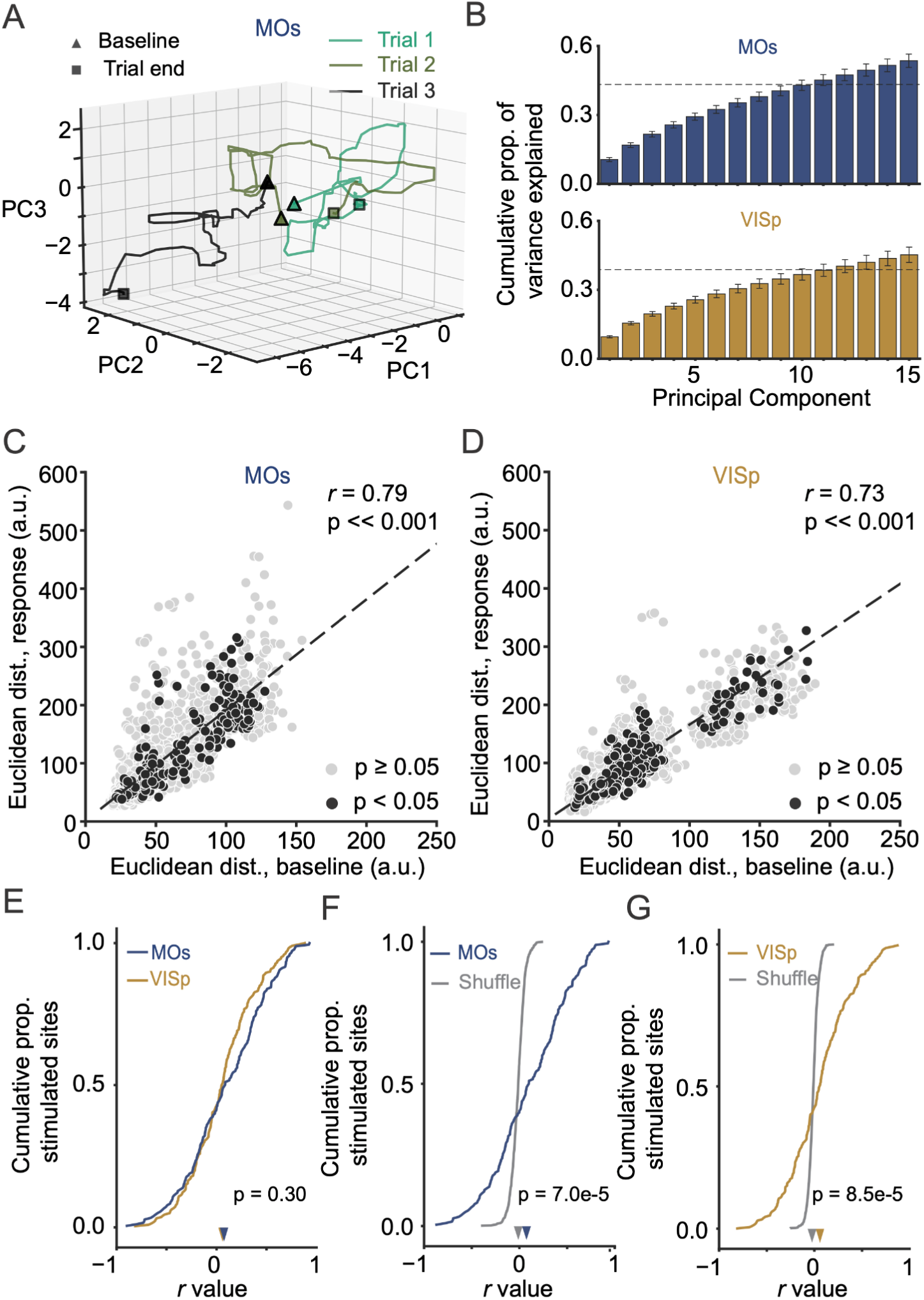
Evoked population-activity trajectories depend on the initial state of the network. **(A)** Single-trial examples of projections onto the first 3 PCs. Note that, compared to the black trace, the two green traces start closer to each other in baseline and show more similar post-stimulation trajectories. Trajectory endpoints are truncated at 3 s post-stimulation for visualization only. **(B)** Cumulative variance explained by the top 15 PCs for each area (VISp: n = 22 FOVs, MOs: n = 18). Bars: ± S.E.M. Dashed line indicates mean variance captured by the top 10 PCs across experiments. **(C)** MOs scatter plots for post-stimulation Euclidean distances between pairs of trial trajectories as a function of distances during the pre-stimulation baseline, computed for trial pairs using the projections onto the top 10 PCs. Black points come from trial pairs for individual photostimulation sites with a significant linear correlation (10 trial pairs each, p < 0.05). The dashed line indicates the linear-regression trend for the aggregate data. Printed *r* and p values are for a linear correlation computed on aggregate data. **(D)** Same as C, for VISp. **(E)** Cumulative distributions of linear correlation coefficient (*r*) as above, across photostimulated sites for both areas. Arrowheads: medians. Printed p-values are from a rank-sum test. **(F)** Comparison of *r* distributions from the MOs data vs. null distributions derived from pseudo-trials obtained with shuffling. Arrowheads: medians. Printed p-values are derived from comparing the data distributions to the median of the shuffled-data distribution. **(G)** Same as F, for VISp.

### Frontal and visual neurons have timescale-dependent response profiles

A diversity of neuronal timescales within a circuit has been proposed to be computationally important, e.g., to provide a flexible reservoir of computation and/or a basis set for temporal integration^5,7,24,69^. However, whether diverse timescales follow a functional connectivity logic, or how they relate to network-level dynamics is still poorly understood. We thus wondered if single-neuron- and network-response patterns to focal photostimulation varied depending on the spontaneous timescales of stimulated and responding sites. We divided the dataset into short- and long-timescale ROIs by separate median splits for MOs and VISp. We then quantified the frequency of significant responses for each of the four pairwise combinations of these timescale categories, within a 10-s time window following photostimulation. Note that, although we lack true cellular resolution for photostimulation, our effective resolution roughly matches the spatial scale of clustering by timescale (**Fig. 2G**). Thus, we are likely primarily targeting sites containing neurons of similar timescales. Moreover, as in our previous analyses, we do not assume this analysis captures short-latency monosynaptic interactions. Interestingly, we found that response probabilities varied according to the timescale of both stimulated and responding sites. In addition to being overall more prevalent in MOs, response probabilities were significantly different depending on timescales in MOs but not VISp (**Fig. 6A**, MOs: χ²_(3, N = 8631)_ = 11.15, p = 0.011; VISp: χ²_(3, N = 13654)_ = 2.21, p = 0.52). Specifically, stimulating long-timescale sites was more likely to yield responses, particularly of other long-timescale ROIs. Next, we used logistic regression to predict whether an ROI would have a significant response to photostimulation, which allowed us to quantify the relative contributions of the timescales of stimulated and responding sites, and of the difference between these two timescales, to response probability (**Fig. 6B**). For both MOs and VISp, the timescale of the photostimulated ROI was significantly predictive of response probability, with longer timescales making it more likely (p < 0.05, subsampling test). Additionally, for MOs, more similar timescales between the stimulated and responding sites significantly predicted higher response probability (i.e., the absolute difference between the two, |Δ*τ*|, p = 0.04). Finally, we asked if the response time courses also depended on timescales. We plotted these time courses for each timescale category, using the same median splits as before. Time courses were very similar in VISp, being always dominated by early responses, but varied depending on timescale for MOs (**Figs. 6C, S4**). Specifically, response peak times happened earlier for short-short and long-long stimulated-responding site pairs (**Fig. 6D**). To quantify this, we fitted a linear regression model to predict peak times from the same features we used in the response probability model above. Indeed, unlike in VISp, earlier MOs peak times were significantly predicted by more similar timescales between the stimulated and responding sites and by longer timescales of the responding neuron (**Fig. 6E**, |Δ*τ*|, p = 0.03 and p = 0.01 respectively, regression t test). This could reflect stronger functional connectivity between neurons with similar timescales. Moreover, combined with the finding of higher response probability between long-long stimulated-responding site pairs in MOs, this arrangement could provide a circuit basis for the selective recruitment of long-timescale neurons during cognitive tasks. This has in fact been observed in the frontal cortex of macaques engaged in a working-memory task^8^.

**Figure 6.**
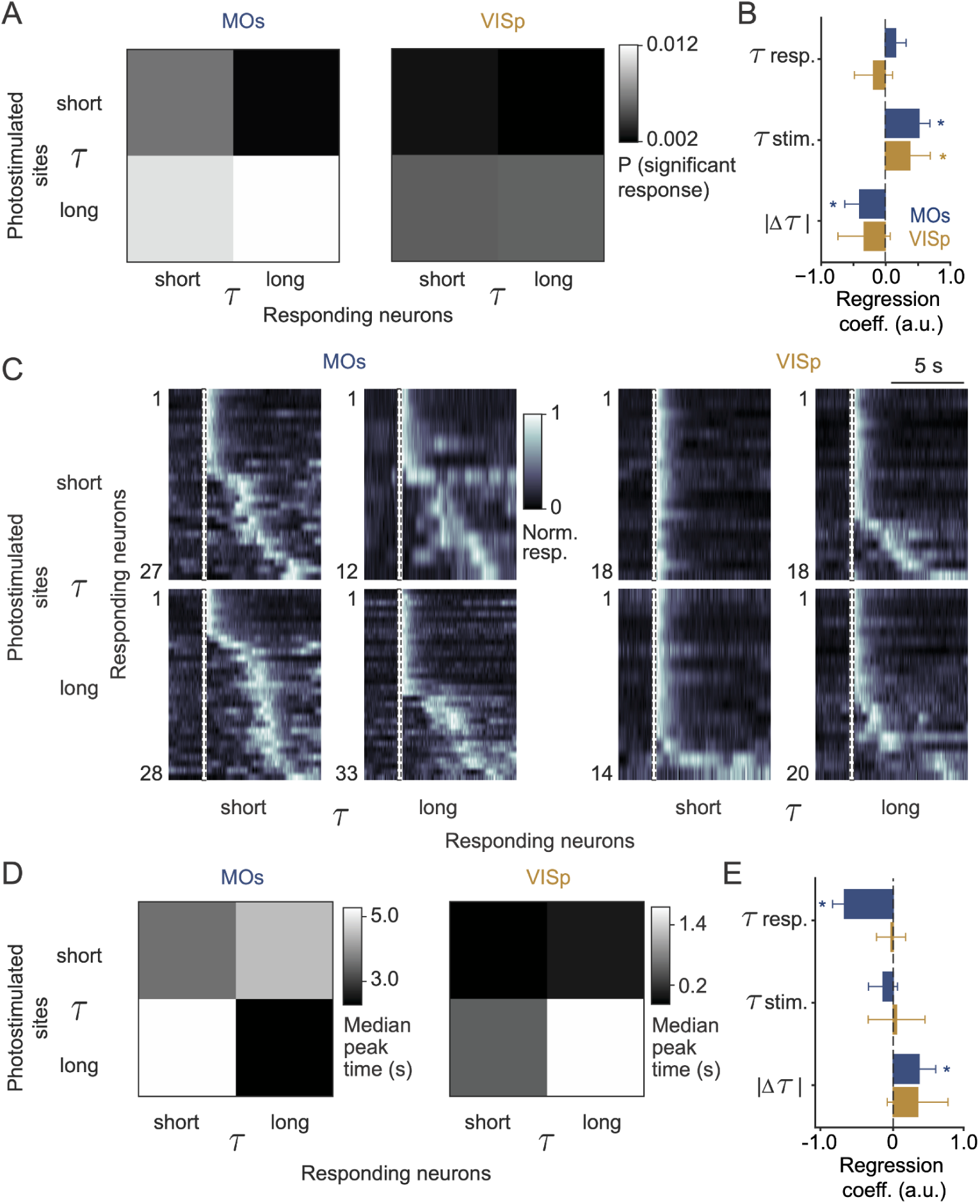
Response probability and time courses depend on timescale. **(A)** Response probability as a function of the spontaneous timescales of the photostimulated and responding sites, categorized by median splits. **(B)** Coefficients from a logistic regression fitted to predict whether responses were significant or not. Bars indicate mean ± S.D. across 1000 fits using different subsamples of non-responding neurons to balance the categories. *: p < 0.05 from the distribution across subsamples. **(C)** Trial-averaged normalized response time courses for all significantly responding neurons in each area that passed criteria for timescale estimation, sorted by peak time and plotted separately for the different timescale-pair categories. **(D)** Median response-peak times for the different categories of timescale pairs. **(E)** Coefficients from a linear regression fitted to predict response peak times. Error bars: standard error of the estimate. *: p < 0.05, regression t test. See also Fig. S4.

## Discussion

Here, we showed that different areas of the mouse cortex have qualitatively distinct types of evoked network dynamics, likely in part due to area-specific circuit-connectivity patterns. First, we developed methods to predict spontaneous cortical timescales from the levels of hundreds of transcripts. We found that, in addition to membrane excitability and excitatory and inhibitory transmission, transcripts associated with circuit wiring are predictive of timescales (**Fig. 1**). Next, using two-photon Ca^2+^ imaging from a visual and a frontal area (VISp and posteromedial MOs), we found that single neurons have broadly distributed timescales, which are slightly longer in MOs, and are spatially clustered by timescales in both regions (**Fig. 2**). Further, we used two-photon methods to deliver brief, focal input to each area while simultaneously measuring evoked dynamics in the network. We found that MOs neurons were significantly more likely than VISp ones to become active upon photostimulation, with timescale-specific response patterns in both areas (**Figs. 3, 6**). However, much like single-neuron timescale distributions, regional differences in response probability were modest. In contrast, the timescales of network-level dynamics were much longer in MOs. This was due to the recruitment of seconds-long activity sequences rather than sustained single-neuron responses (**Fig. 4**). Finally, these network-level evoked dynamics were partially determined by the state of the network at the time of stimulation (**Fig. 5**). Overall, our results are compatible with the idea that longer timescales in frontal cortical regions arise in part from network-level interactions between neurons, enabled by stronger and/or more prevalent synaptic connectivity. To our knowledge, this work provides the first direct comparison of cortical-area-specific network dynamics using causal perturbations *in vivo*. Our findings have important implications for a mechanistic understanding of how different areas take part in cognitive processes like decision making and working memory.

Previous efforts to quantify cytoarchitectonic and transcriptomic features related to timescales primarily relied on differences between cortical areas, which in turn differ in their timescales^21–23^. We built on these efforts to directly predict timescales from transcript levels in a way that is agnostic to anatomical location and confined to a single known neuronal population. This approach allowed us to identify novel transcripts that could be involved in setting circuit timescales by influencing circuit wiring. This agrees with studies of dendritic morphology that suggest stronger excitatory connectivity in frontal areas^19,21^, and identifies possible molecular targets for future studies of what mechanisms regulate area-specific connectivity.

Our transcriptomics analysis also highlights the fact that multiple biological phenomena, from the sub-cellular to the network levels, are likely to collectively give rise to cortical timescales. On the other hand, our functional data show that network-level differences in evoked cortical dynamics are much stronger than differences in single-neuron firing properties. Further, because the dynamics we observed in frontal area MOs evolve in the form of seconds-long sequences, we conclude that it emerges from (presumably polysynaptic) recurrent interactions rather than cell-autonomous mechanisms. Note, however, that our approach does not allow us to differentiate between local and long-range excitatory connectivity. Indeed, the latter has been shown to be important for long-timescale dynamics in both computational models and mice performing working-memory tasks^30,70–73^. Future work should dissect the relative contributions of connections within vs. across areas for network dynamics. Likewise, although we observed similar levels of inhibition following optogenetic stimulation in both regions, future studies should investigate the potential role of area-specific inhibitory mechanisms in shaping their dynamics, since the relative proportions of inhibitory-neuron subtypes also changes along the cortical hierarchy^21,22^. Specifically, VISp has relatively more parvalbumin-positive interneurons than MOs. This inhibitory subtype provides potent perisomatic inhibition^74^, which could counteract recurrent excitation to shorten network responses. Finally, potential differences in short-term synaptic plasticity between regions could also play a role — longer-lasting dynamics could also take place if facilitation is more common in frontal than sensory areas (e.g., ^75^ vs. ^76^). However, we are unaware of any systematic comparisons of this property across the cortex.

Regardless of the exact underlying mechanisms, our data suggest that frontal circuits are intrinsically better equipped than visual ones to generate long-lasting dynamics via activity sequences, given that our experiments were done in non-task-trained mice. This could help explain why these areas are differentially engaged in cognitive processes — for instance, mouse MOs seems to be primarily recruited in working-memory tasks during long memory delays^14,77^. Note, however, that activity sequences have been observed across multiple visual, frontal and subcortical areas^57–65^, and that we did observe late-responding cells in VISp, albeit in much smaller proportion than MOs. Moreover, activity timescales change with training and are modulated by behavioral-task performance in rodents and primates^3,13,43,78^. This suggests that longer-lasting network dynamics can be acquired with training. An open question is whether these trained dynamics are enabled by longer sequences or persistent activity at the single-unit level. Our data from the cortex of naïve mice provide little evidence for prevalent sustained responses, perhaps suggesting that this is acquired with training. The exact answer likely depends on exact task design, brain region and species (e.g., ^8,25,27,33,43,57,59,79–81^).

Thus, much work remains before we have a complete understanding of how distinct cortical circuits can perform computations on different timescales via network dynamics, and how their intrinsic properties can be modified with learning to enable cognitive processing. Our results provide an important step forward. Beyond the direct demonstration of distinct dynamics, they should help constrain computational models of cortical function that explicitly take into account differences between cortical circuits.

## Methods

### Experimental Animals

We used male and female Thy1-GCaMP6s mice aged 12–60 weeks at the time of data collection (n = 10, line GP4.3^82^, Jax stock # 024275). Animals were housed in a temperature-controlled environment with a reverse 12-hour light/dark cycle and provided with food and water ad libitum. All procedures were approved by Northwestern University’s Institutional Animal Care and Use Committee (IACUC) and conducted in accordance with the National Institutes of Health Guide for the Care and Use of Laboratory Animals. Widefield Ca^2+^ imaging data were reanalyzed from a previous publication and collected from Cux2-Cre x Ai96 crosses as detailed previously (n = 5)^14^.

### Surgical Preparation

Mice aged 8–12 weeks underwent a single procedure to virally deliver ChRmine and implant a circular glass window over either VISp or MOs (5-mm diameter, #0 coverlip). They were anesthetized using isoflurane (3% induction, 1.5–2.5% maintenance) and placed in a stereotaxic frame (Kopf). Scalp hair was removed using Nair^TM^ and the scalp was disinfected with betadine and 70% ethanol. Lidocaine (4% w/v) was applied subcutaneously for local anesthesia. The skull was exposed using a large incision and a cranial window (∼5 mm diameter) was created using a dental drill and fine forceps. For MOs, the window was centered at bregma, and for VISp, at +1.0 mm AP and 2.0 mm ML from lambda. We then used a microinjection syringe pump (World Precision Instruments) to deliver AAV8-CaMKIIa-ChRmine-mScarlet 500 µm below the pial surface (1-3 injections per mouse, 500 nL, rate of 50-100 nl/min, titer ∼10^12^). The syringe was retracted 5–10 min after the injection. Injection coordinates for MOs were AP +1.0 mm and ML 0.5–0.75 mm from bregma and, for VISp, AP +1.0 mm and ML 2.0–2.5 mm from lambda. The circular glass window was then lowered onto the craniotomy using a manipulator and bonded to the skull using veterinary cyanoacrylate glue and metabond (Parkell). Finally, a titanium headplate was bonded to the skull using opaque metabond mixed with carbon powder. Body temperature was maintained at 37°C throughout the procedure using a homeothermic blanket (Harvard). To reduce brain swelling, dexamethasone was administered via intramuscular injection (2 mg/kg) 1 hour before the beginning of the surgery. For analgesia, mice received a subcutaneous injection of meloxicam (20 mg/kg) before the procedure and 24h later, as well a sustained-release formulation of buprenorphine (Buprenorphine SR, 1 mg/kg) subcutaneously before the procedure for prolonged pain relief, lasting up to 72 hours. They also received warm saline after the procedure to maintain hydration (0.1–0.5 mL S.C). The mice were allowed to recover for at least 7 days before being habituated to handling, followed by habituation to head fixation by progressively increasing its duration from 10 to 90 min in 15–30-min increments.

### Spatial transcriptomics analyses

#### Datasets

We used the functional spontaneous-timescales widefield imaging dataset from ^14^ (54-µm/pixel resolution, see above for mice and below for timescale estimation methods). The transcriptomics dataset was a publicly available MERFISH-imputed spatial transcripts dataset from ^38^, downloaded via the Allen ABC Atlas Access API. This dataset includes imputed values of transcript expression for 8,460 unique transcripts with a spatial resolution of 10 µm. To preserve statistical power in our regression, we hand-selected a subset of 252 transcripts for several families of ion channels and ion homeostasis-related proteins, neurotransmitter receptor subunits and transporters, synaptic and adhesion proteins, transcription factors and proteins implicated in plasticity and development, transcripts previously used in classifying cortical neuronal subtypes^39^, and negative-control transcripts involved in non-neuronal-specific metabolic function (full list in **Tables S1, S2**).

#### Dataset co-registration, data selection and resampling

We used bregma and lambda as fiducial markers for registering each mouse in our functional dataset to the Allen ccf3.0 using an affine transformation with bregma at [−5.2 mm AP, 5.7 mm ML] and lambda at [−9.3 mm AP, 5.7 mm ML]. MERFISH-imputed data are already provided in ccf3.0. For simplicity, coordinates were converted to mm relative to ccf origin for both datasets, with the AP axis being inverted such that more positive values corresponded to anterior cortical regions. We excluded widefield pixels with signal-to-noise ratio (SNR) < 3 and whose exponential fit for *τ* estimation had an *R*^2^ < 0.8^14^. Because all the widefield data was collected from Cux2-Cre mice, we only included transcript-level data from neurons with non-zero levels of *Cux2*. The exception was the control analysis in Fig. 1D, in which we restricted transcripts to layer-6 excitatory neurons found in the same AP-ML coordinate as the widefield pixels. We pooled both datasets into isotropic 100-µm voxels (pixels), and resampled expression levels and functional timescales within each voxel, with replacement, as follows. For each widefield pixel within a registered voxel, we randomly sampled expression levels once. The total number of widefield pixels in a voxel thus determined resampling counts for each voxel, such that the functional dataset was the bottleneck for the co-registration process, to prevent over-representation of transcriptomic profiles where functional data was lacking. This was done over the aggregate of all sessions, and voxels with fewer than three observed MERFISH-imputed neurons or widefield-imaging pixels were excluded. Finally, we made the analysis hemisphere-agnostic by collapsing all voxels along the midline for all analysis by centering the sagittal midline on zero and taking the absolute value of the ML coordinates.

#### Transcript-enrichment analysis

For the analysis in Fig. 1C, we calculated the average copy numbers of each transcript across co-registered voxels belonging to the top and bottom quartiles of the *τ* distribution computed across resampled data. We then computed the log_2_ of the ratio between long- and short-timescale pixels. We computed p-values for each transcript with a two-sided t test between the pixel-wise copy-number distributions within the two timescale quartiles, with false discovery rate (FDR) correction performed as described below (*General statistics*).

#### Linear regression models

For the analyses in **Figs. 1D, E** and **S1** we used L1-regularized linear regression with the transcript levels and/or co-registered ML and AP position as predictors, *τ* as the response variable, and resampled pixels as observations. All predictor and response-variable columns were z-scored independently. For regularization, we used an 80-20 training-testing split and inner-outer cross-validation. We used an inner 5-fold cross-validation to select the L1 regularization term, *α*(*α* = 10^-5^ – 10^-2^ in 30 equally sized steps), which was then used to fit the model on the full training set, with final model predictions made on the test set. The outer test cross-validation was done over 5 folds for a total of 5 fitted models per regression type. Reported model coefficients and goodness of fit (*R*^2^) were calculated as the mean ± S.D. across these 5 outer folds. P-values for model coefficients were computed using a two-sided t test against zero, with the t statistic being computed as the mean coefficient divided by its standard error with degrees of freedom equal to 4 (N folds minus 1). The p-values were then FDR corrected. The model in Fig. 1E only had the 252 transcript as predictors. The model in **Fig. S1A** additionally had ML and AP positions as predictors. Finally, for the analysis in **Fig. S1B**, we fitted a model as in Fig. 1E and another one with ML and AP positions as a two-dimensional response variable and the 252 transcripts as predictors. We then defined transcripts that were predictive of *τ* but not AP anatomical position as those with coefficients outside the central 10th-90th percentile of the coefficient distribution for the timescales model, but within the central 10th-90th percentile AP coefficients for the spatial model.

### Behavioral apparatus and experimental design

Experiments happened while head-fixed mice ran spontaneously in a dark environment. They sat on an 8” hollow Styrofoam© ball suspended by compressed air. We measured running speed at 120 Hz via ball-surface displacements using an optical velocity sensor (ADNS-3080 APM2.6) housed in a custom 3D-printed cup. The sensor was positioned underneath the ball and was controlled by an Arduino Due connected to a PC running the Matlab-based virtual reality engine ViRMEn. Each experiment consisted of a structural z-stack to characterize ChRmine expression, immediately followed by a 5–10-min baseline imaging session for the estimation of spontaneous activity timescales, and then one of four phostimulation protocols as described below (duration ∼50–80 min).

### Two-Photon Ca^2+^ Imaging

Two-photon microscopy was performed using an Ultima 2Pplus two-photon laser scanning microscope (Bruker) equipped with mode-locked fiber lasers with acousto-optic power modulators (Coherent Axons, 80-MHz repetition rate, 920 and 1064 nm for GCaMP6s and mScarlett imaging, respectively). Laser beams were combined using a dichroric and scanned at 30 Hz using an 8-kHz resonant galvanometer over a ∼500 × 500-µm field of view (FOV, resolution of 1.1 µm/pixel). An electrotunable lens (ETL) in the imaging path allowed us to move the imaging plane along the axial (z) direction independently of the mechanical movement of the objective. Emitted fluorescence was collected through a 16× water-immersion objective (Nikon, NA 0.8), passed through emission filters (et525/50m-2p and et595/50m-2p) and detected using Galium Arsenide Phosphide photomultiplier tubes (GaAsP PMTs, Hamamatsu, H10770PB-40). Based on preliminary validation experiments (not shown), power for the 920-nm laser was maintained below 40 mW to avoid cross-spectral activation of ChRmine. All time series datasets were acquired from layer 2/3 at a depth of 140–200 µm from the pial surface. MOs FOVs were centered around AP +0.5–1.0 mm and ML 0.5–1.0 mm from bregma, and from VISp around AP +1.0 mm and ML 2.0–2.5 mm from lambda. Two-photon data were acquired using PrarieView (Bruker), and synchronized with behavioral running data by acquiring a copy of the galvanometer command voltage directly onto the behavioral data acquisition software running on a separate PC. We collected data separately from mice implanted with cranial windows in MOs (n = 5 mice, 64 FOVs) or VISp (n = 5 mice, 69 FOVs).

### Focal two-photon optogenetic stimulation

We used a separate pathway in the microscope for focal two-photon optogenetic activation. ChRmine excitation was done with a 1035-nm laser with 1-MHz repetition rate and 250-fs pulse width (Coherent Monaco), combined with the imaging laser right before the objective using a 1040 notch-filter dichroic (1040drcb). Phostimulation sites were manually selected based on verified ChRmine expression with a 1064-nm-excitation morphological scan for mScarlet excitation. Selection was blind to the functional properties of the neurons. The laser beam (5 mW) was centered on the somata and spiraled outward using x-y line-scanning galvanometers (spiral diameter: 15 µm). Photostimulation patterns were generated and controlled using the MarkPoints feature of the PrairieView software. The PMTs were shuttered during photostimulation.

#### Stimulation protocols

In most experiments (**Figs. 3, 4A–F, 5, 6, S2C–E, S3D–E, S4**, n = 9 mice, 40 FOVs, 400 stimulation sites), the photostimulus consisted of 5 spirals (15 ms on, 35 ms off, total 250 ms). We stimulated 10 individual sites in pseudorandom order, with a 30-s inter-stimulus interval (ISI). Each site was stimulated 5 times, with a minimum interval of 300 s between trials. For the experiments in **Figs. S2F, G**, we delivered a single 15-ms spiral, with the same overall design as above (n = 5 mice, 24 FOVs, 240 stimulation sites). For the experiments in **Figs. 4G, H**, we selected a single site and stimulated it 20 times with an ISI of 180 s (n = 2 mice, 12 FOVs, 12 stimulation sites). Finally, for the control experiments measuring the spatial resolution of two-photon optogenetics (**Fig. S2A, B**, n = 5 mice, 48 FOVs, 48 stimulation sites), we selected a single layer-2/3 neuron expressing ChRmine and moved the stimulation sites away from that neuron, with 5 trials for each distance step. For experiments testing lateral resolution, the site was moved in 7.5 µm steps. For experiments testing axial resolution, to ensure that we continued imaging the same focal plane while changing the position of the stimulation site in the z-axis, we introduced a z-offset between the stimulation plane and the imaging plane. We used two independent devices for axial control: an ETL integrated into the imaging path, and a z-axis stage motor that physically moved the objective. We systematically shifted the stimulation site along the z-axis in 20 µm steps, while keeping the imaging plane fixed on the same focal layer.

### Two-photon data preprocessing

Time series data were preprocessed using Suite2p^83^. Briefly, imaging frames were corrected for lateral motion by maximizing the phase correlation with a reference average frame. Regions of interest (ROIs) were then segmented using the parameters below, followed by semi-automated curation to extract only ROIs that corresponded to neuronal somata. Raw fluorescence for each ROI was then extracted as the average fluorescence of all pixels assigned to it. For the analysis of spontaneous activity in Fig. 2, 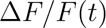 was calculated as 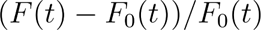, where *F*_0_ was defined as the mode of *F* over a 180-s running window. Photostimulation-triggered activity was computed directly on raw fluorescence traces as explained below (*Analysis of neuronal responses to optogenetic stimulation*).

### Suite2p parameters

| Parameter | Value | Parameter | Value |
| --- | --- | --- | --- |
| suite2p_version | 0.10.3 | subpixel | 10 |
| smooth_sigma | 1.15 | snr_thresh | 1.2 |
| spatial_hp_reg | 42.0 | spatial_scale | 2 |
| max_iterations | 20 | threshold_scaling | 0.75 |
| max_overlap | 0.75 | pretrained_model | cyto |
| soma_crop | 1.0 | min_neuropil_pixels | 350 |

### Estimation of spontaneous-activity timescales

We estimated timescales following the method we have recently described and validated using simulations^14^. Briefly, we first removed the direct contributions of running to the activity of each ROI by fitting the following L2-regularized linear model with running speed at multiple lags as predictors:

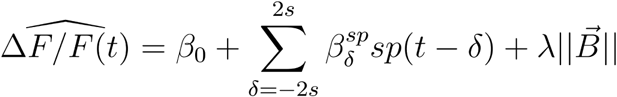

Where *β*’s are the model coefficients, *δ*’s are the lags, *sp_t_* is running speed at time point *t*, ||*B⃗*|| is the L2-norm of the coefficient vector and *λ* is the regularization parameter, determined by cross-validation. We then extracted the residuals of this regression and computed their autocorrelation function with a maximum lag of 30 s for two-photon data and 60 s for widefield data. The values at non-negative lags of the autocorrelation function *G*(*δ*) were then fitted with both a single and a double exponential, 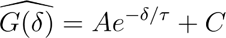 and 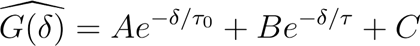. *τ*_0_ was bounded between 0 and the Nyquist period of the signal (i.e., ∼60 ms), and *τ* was bounded between the Nyquist period and 100 s. We then selected the fit with the highest *R*^2^ and defined the spontaneous timescale as the *τ* term from the best fitting exponential.

### Data selection

For the analyses of single-neuron timescales, we excluded ROIs with a best-fitting *R*^2^ < 0.8, signal-to-noise ratio (SNR) < 2, and fewer than one significant transient per min. Based on the intuition that most signal in Ca^2+^ imaging data lies in low frequencies, we defined 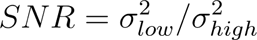, where *low* is ΔF/F low-pass filtered at 9.9 Hz, *high* is ΔF/F high-pass filtered at 10.1 Hz (20th-order finite impulse response filters in both cases), and *σ*^2^ is the variance of the filtered traces. Finally, we estimated significant transients as follows. We first estimated a noise distribution as the absolute value of all negative transients, and set a significant threshold for positive transients as three standard deviations of the noise distribution. We considered positive transients to be significant if they exceed this threshold for at least a consecutive 200 ms and if at least 80% of data points within the whole transient were above this threshold. Using these criteria, we selected 2,387/6,480 VISp and 1,501/5,808 MOs ROIs for further analysis in **Figs. 2, 6** and **S4**. For the analysis of responses to photostimulation (**Figs. 3–5, S2–S3**), we excluded all ROIs with centroids within a 25-µm radius of the center of the phostimulated site but did not apply SNR or event-rate criteria under the assumption that even silent ROIs could respond to photostimulation.

### Analysis of spatial clustering by timescale

For the analysis in Fig. 2G, we z-scored *τ* values, and computed all unique pairwise spatial distances between the centroids of each ROI and the absolute differences between their *τ* as computed above, |Δ*τ*|, all within each FOV. We then concatenated all the data into a single vector of distances and one of |Δ*τ*|. We bootstrapped the concatenated data for 10,000 iterations to compute distributions of |Δ*τ*| per binned spatial distances. Additionally, we shuffled the distance vectors 1,000 times to estimate empirical chance distributions of |Δ*τ*| as a function of space. The values in Fig. 2G were obtained by subtracting the shuffled values from the bootstrapped distributions, such that they reflect how much closer timescales are than expected by chance. Two-sided p-values were calculated as the proportion of shuffled-subtracted bootstrapping iterations with a sign opposite to the median one.

### Analysis of neuronal responses to optogenetic stimulation

Raw fluorescence traces were aligned to the onset of phostimulation separately for each uniquely targeted site, and timepoints during PMT shuttering were assigned NaN values. Each single-trial trace was then smoothed using a ∼200-ms (7-timepoint) median filter. To standardize responses across cells and sessions, single-trial smoothed traces were z-scored to a 3-s pre-stimulus baseline. We identified peak changes in fluorescence (times and width) using the ‘find_peaks’ function from the SciPy toolkit, which detects local maxima by comparing each point to its neighbors; the width parameter within ‘find_peaks’ function was set to 2 points. Neurons were deemed to respond significantly to photostimulation if they passed all the following criteria: 1) their average peak response exceeded a z-score of 3 for at least 2 consecutive timepoints; 2) they had reliable evoked responses, defined as at least 60% of trials having peak transients above 3 for at least 2 consecutive timepoints; 3) their peak responses exceeding the threshold occurred at consistent times after photostimulation onset, defined as having a standard deviation across peak times smaller than 0.75 s. For analyses of peak widths, peak times, and peak amplitudes of responding cells (**Fig. 3G–I, Fig. 4E–F, Fig. S2E–G**) we only included values from trials with peak amplitude larger than our z-score cutoff listed above. Response probabilities in Fig. 3G were computed separately for each experiment. Finally, in **Fig. S2E**, we defined significant inhibition by photostimulation as the following criteria: 1) their average response decreased by at least a z-score of 2 for at least 2 consecutive timepoints; 2) they had reliable evoked responses, defined as at least 60% of trials having decreased below 1 z-score for at least 2 consecutive timepoints.

#### Estimation of population-response timescales

For the analysis in Fig. 4A, we averaged the peak-normalized, trial-averaged responses of all significantly responding neurons as defined above. Note that the same neuron can contribute to the data more than once if it responds significantly to more than one stimulated site within the same FOV (the same is true for Fig. 4E–G). To estimate the timescales of evoked responses in Fig. 4B, C, we selected random subsets of *n* significantly responding neurons, with n = 1, 5, 10, 25, 50, or 100. For each *n*, we drew 1,000 random subsets, averaged their responses as above, and computed the autocorrelation function of the average response. Computing the autocorrelation instead of directly using triggered averages allowed us to capture occasional non-monotonic responses (e.g. from the selection of large subsets of late responding cells). We then fitted a single or double exponential decay function to the autocorrelation to extract a single *τ* as explained above. The p-values comparing the *τ* distributions of VISp and MOs were computed as the proportion of random subsamples for one area that crossed the median of the subsamples of the other area.

#### Estimation of null peak-time distributions

For the analysis in **Fig. S3A, B**, in addition to the significance criteria above, we devised a more stringent procedure to estimate the false positive rate arising from spontaneous fluctuations in activity. For each significantly responding neuron, we generated pseudo-trials by segmenting non-overlapping time windows from the 5–10-min initial session used to estimate spontaneous timescales, to preserve cell-specific general activity levels. These trials were matched in duration and number to the actual photostimulation-triggered traces and underwent identical z-scoring and data selection procedures. This was repeated 1,000 times, and a neuron was deemed to still be significant if its pseudo-trial controls were selected as a significant response fewer than 5% of times.

#### Cross-validation of peak times

For the analysis in **Fig. 4G, H**, we devised a cross-validation procedure to estimate the consistency of peak times across different instances of photostimulation for the dataset with 20 trials/site. Activated neurons were only selected based on significance criteria 1 and 2, to avoid pre-selection by peak-time consistency. The 20 trials were randomly into halves 10 times, and median peak times across the 10 repeats were extracted for each half. A linear correlation coefficient *r* between trial halves was then computed across neurons. This procedure was repeated 1,000 times to generate an *r* distribution, which was compared to the distribution obtained from pseudo-trials constructed as described above. Statistical significance between the actual and pseudo-trial distributions was computed as the proportion of trial-half *r* values that crossed the median of the pseudo-trial distribution. For the analysis in **Fig. S3D, E**, we repeated this same procedure for data collected using 5 trial repeats instead of 20 trials.

### Photostimulation-triggered running speed

Running speed was defined as the Euclidean norm of the time derivative of the x-y displacement recorded with the optical velocity sensor. Speed traces were then aligned to individual photostimulation trials and z-scored to a 3-s baseline as done for neuronal responses. An average photostimulation-triggered trace was then computed for each unique stimulated site and averaged across sites for the analysis in **Fig. S3C**.

### Quantification of ChRmine expression strength

For the analysis in **Fig. S2C, D**, because the ChRmine construct is fused to mScarlet, we extracted red fluorescence values as a proxy of expression strength. We applied the Suite2P ROI masks to the red channel of the relevant depth within the morphological z scan acquired before the time series, and averaged all pixels within the ROI. To normalize expression levels across FOVs, we z-scored red fluorescence to the distribution obtained from all segmented somata, separately within each FOV.

### Principal Components Analysis (PCA)

To assess the relationship between baseline activity and trial-to-trial variability at the population level (**Fig. 5**), we performed PCA on the activity of all imaged neurons within an FOV, regardless of whether they were deemed as significantly responding to the photostimulus. For each experiment, we first constructed a population-activity matrix in which each row represented a timepoint and each column corresponded to a neuron’s z-scored ΔF/F measured during the 5–10-min initial session used to estimate spontaneous timescales. We then decomposed this matrix using the randomized singular value decomposition method (scikit-learn implementation). We separately projected the activity of each photostimulation trial onto the space spanned by the top *K* PCs, with *K* = 3 for visualization and 10 for quantification. Euclidean distances were computed across all timepoints for the 3 s preceding photostimulation (baseline) or the 7.5 s following it (response). Using different values of *K* or durations of baseline and response periods did not qualitatively change any of the findings (not shown).

### Analysis of photostimulation-evoked activity by spontaneous timescale

For the analyses in **Fig. 6**, cells were split according to the median spontaneous timescale (with the inclusion criteria applied as above), with single medians computed separately for each region across all experiments. Response probabilities in **Fig. 6A** were computed separately for each experiment and then averaged (for consistency with **Fig. 3G**). If ROIs of a given timescale category did not exist in a given experiment (which is not guaranteed due to the single median), response probabilities were assigned a NaN value and not factored into the average. Using an overall probability instead of single-experiment averages yielded similar results (not shown).

#### Logistic regression of response probability

In **Fig. 6B**, we fitted a logistic regression to predict the probability of a significant response to photostimulation, *P_resp_*

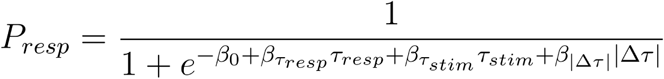

where *τ_resp_* is the spontaneous timescale of the responding neuron, *τ_stim_* is the spontaneous timescale of the stimulated site, and |Δ*τ*| is the absolute difference between *τ_resp_* and *τ_stim_*. Because of the very low overall response probability, we devised a resampling procedure to balance the numbers of responding and non-responding neurons and maintain statistical power. In each of 1,000 iterations, we kept all the *N* responding neurons and sampled a random subset of *N* non-responding ones without replacement and fitted the model above. In the figure, we report the mean and standard deviation of fitted coefficients across iterations. P-values within regions were computed as the proportion of iterations with coefficients of sign opposite the sign of the mean value, and across regions as the proportion of the per-iteration difference between coefficients that were below (above) zero, depending on the sign of the mean difference.

#### Linear regression of peak times

In **Fig. 6E**, we fitted an ordinary least squares (OLS) linear regression model to the peak times of significantly responding neurons, using the same predictors as in **Fig. 6B**. In this case we ran a single model and extracted error estimates and p-values from model residuals, and t statistics using standard regression methods.

### General statistics

Shuffle-, resampling-, and bootstrapping-based tests are detailed in their corresponding sections. For all other analyses, distributions were first assessed for normality using the Shapiro-Wilk test. We then used t-tests for normally distributed data and rank-sum tests otherwise. For comparisons between multiple groups in **Fig. 3J**, we applied a two-way analysis of variance (ANOVA). We used FDR correction^84^ for multiple comparisons (e.g., transcriptomics analysis, spatial clustering analysis).

## Acknowledgements

This work was supported by the US National Institutes of Health (NIH) grants R00MH120047 (to L.P.), R01MH138285 (to L.P.); Simons Foundation grants 872599SPI (to L.P.), NC-GB-PilotExt-00002091 (to L.P.); Alfred P. Sloan Foundation grant SP-2022-19027 (to L.P.); Scialog grant 29059 from the Research Corporation for Science Advancement (to L.P.); T32 (to J.E.C. NIH parent award T32 AG020506); NURTURE postdoctoral fellowship (NIH parent award U54CA272163) (to R.M.C.). We thank Xiaoyin Chen, Andrew Miri, Joshua Glaser and James Fitzgerald for helpful suggestions on the transcriptomics analysis, and Julia Cox and Gordon Shepherd for thoughtful comments on this manuscript.

## Author contributions

L.P. and J.E.C. designed the research. R.M.C. collected the widefield data. J.E.C. performed the two-photon microscopy experiments. J.E.C. analyzed the two-photon data. L.A.A.-S. performed the transcriptomics analysis. L.P., J.E.C. and R.M.C. secured the funding. L.P. and J.E.C. wrote the paper. L.P. supervised the work.

## Data and code availability

The full dataset and source code will be publicly released upon publication of this work.

## Declaration of interests

The authors declare no competing interests.

## Supplemental Information

**Figure S1.**
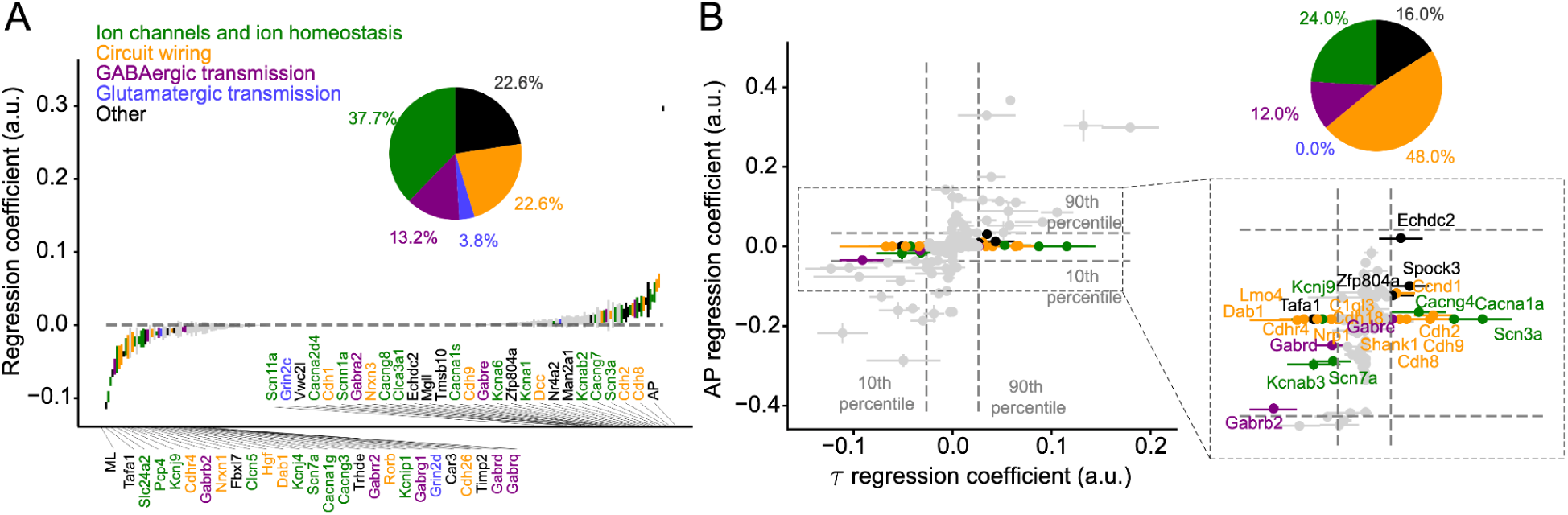
Transcripts implicated in circuit wiring remain predictive of timescales when explicitly accounting for their anatomical position. **(A)** Regression coefficients from the L1-regularized model that predicts pixel-level timescale from transcripts in layer-2/3 *Cux2*-containing neurons, as well as the AP and ML position of the pixel. Only significant predictors are labeled, as defined by p < 0.001 with FDR correction (derived from a two-sided t test against zero), and color coded according to the categories in the legend. Inset: pie chart showing the proportion of significant transcripts in each manually defined category. **(B)** Scatter plot of regression coefficients (transcripts) from the *τ* model in Fig. 1E (x axis) vs. the AP-regression coefficients from a model fit to jointly predict the AP and ML position of a pixel (y axis). Only significant predictors are labeled (inset), as defined by transcripts that are beyond (below) the 90th (10th) percentile of the coefficient-magnitude distribution for the *τ* but not the AP/ML model. Pie chart conventions as in (A).

**Figure S2.**
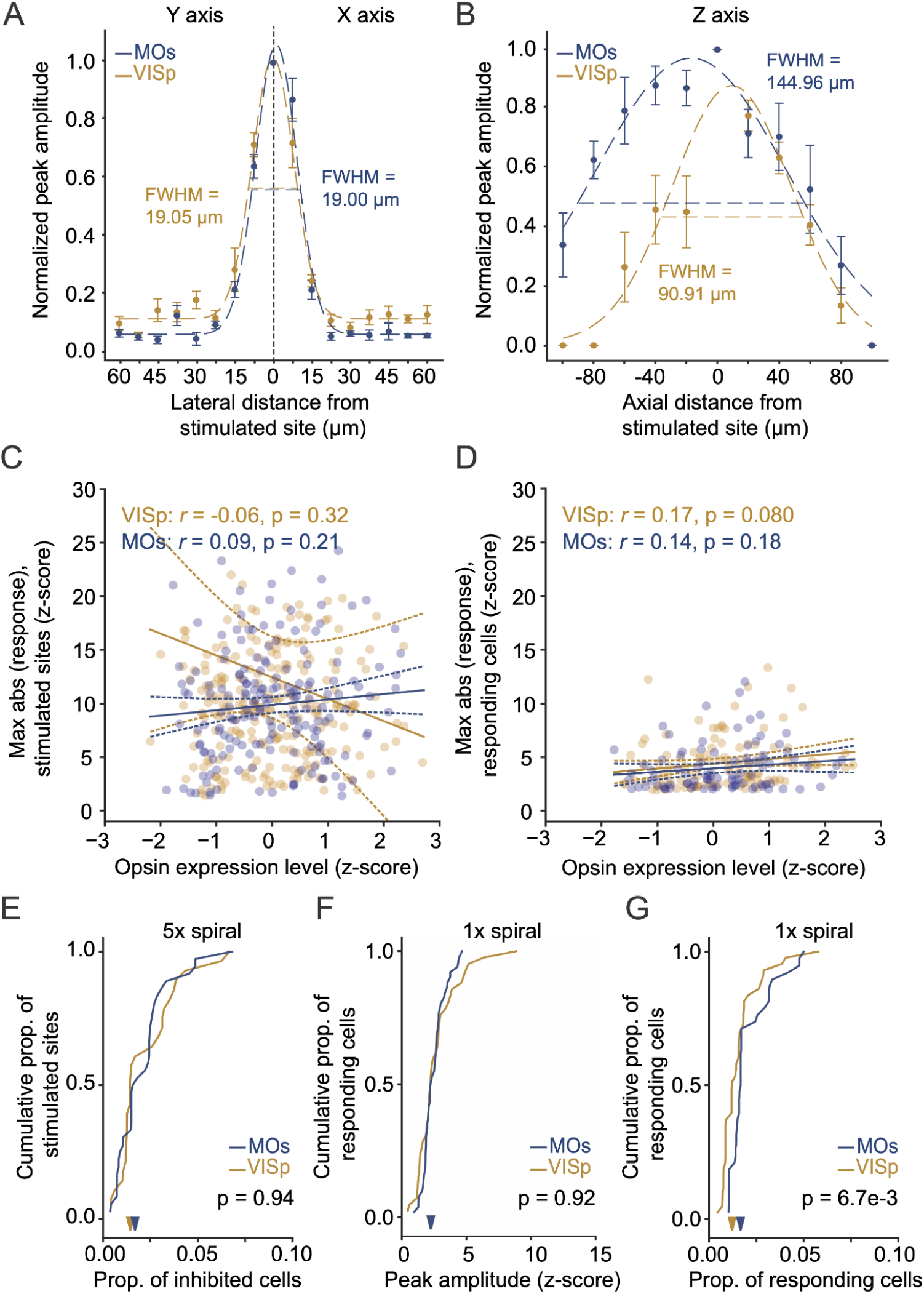
Additional findings and characterization of focal photostimulation. **(A)** Lateral profiles of responses (x and y axes). Normalized peak amplitudes are plotted as a function of distance from the stimulation site for neurons in MOs and VISp. Error bars: ± S.E.M. Dotted lines: best-fitting Gaussian curves, with full width at half maximum (FWHM) values indicated. **(B)** Same as (A), along the axial (z) direction. **(C)** Relationship between opsin expression level (z-score) and maximum response amplitude (z-score) for stimulated sites. Data points: ROIs, solid lines: linear regression trends, dashed lines: linear regression 95% confidence intervals. Printed *r* and p-values are from linear correlation. **(D)** Relationship between opsin expression level (z-score) of the stimulated site and maximum response amplitude (z-score) of responding sites. Conventions as in (C). **(E)** Cumulative proportion of significantly inhibited cells in each region across stimulated sites using a 5-spiral stimulus (same as the one used in Figs. 3–6). Arrowheads: medians. Printed p-value is from a rank-sum test. **(F)** Cumulative distribution of peak-response magnitude from directly stimulated sites in VISp (n = 54from 9 FOVs in 3 mice) and MOs (n = 90 from 9 FOVs in 3 mice), for experiments using a single spiral. Arrowheads: medians. Printed p-value is from a rank-sum test. **(G)** Cumulative distribution of the proportion of significantly responsive neurons for the photostimulated sites in (F) (i.e., single spirals). Arrowheads: medians. Printed p-value is from a rank-sum test.

**Figure S3.**
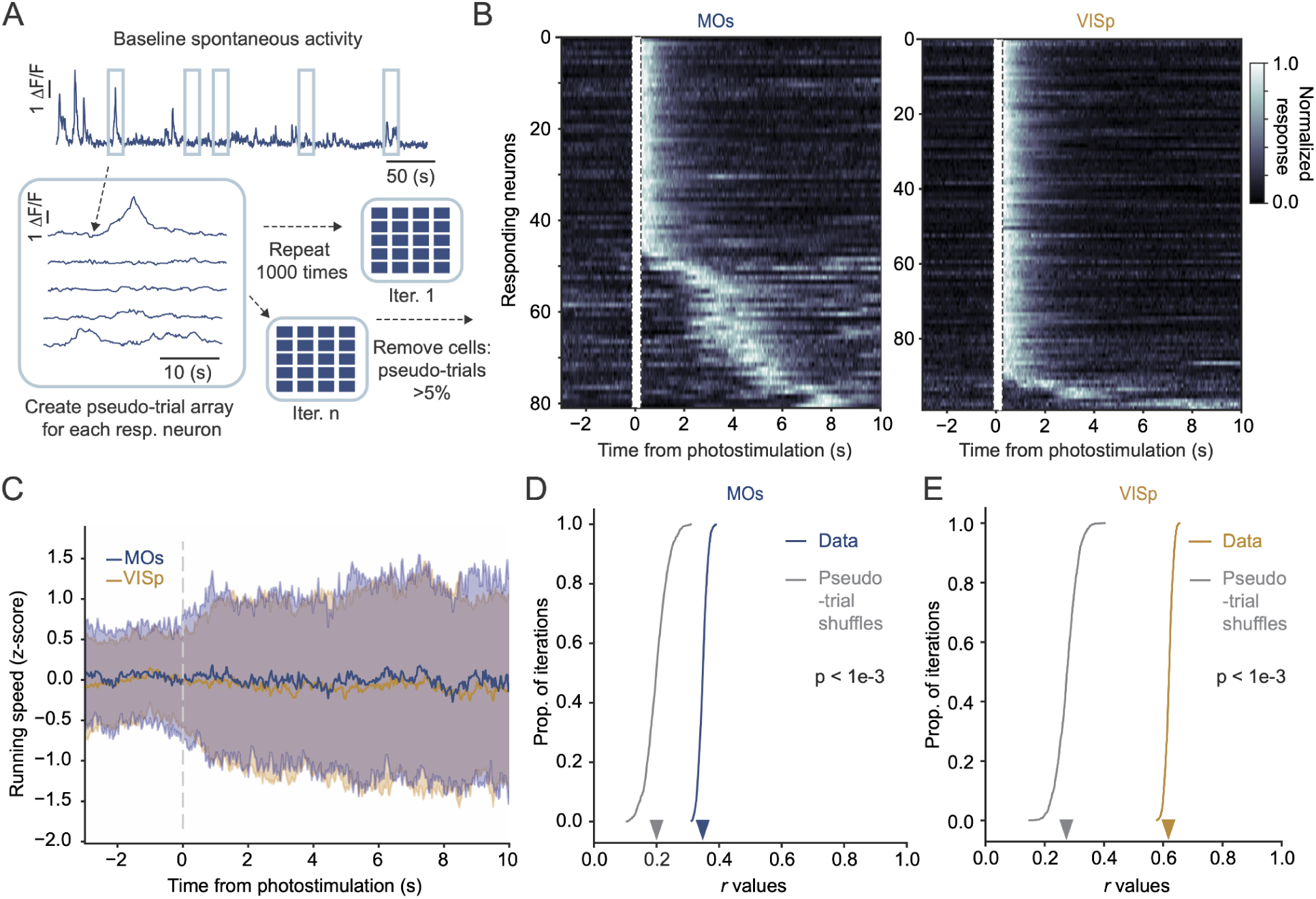
Further analysis of evoked network dynamics. **(A)** Schematic of pseudo-trial generation from spontaneous activity. For each significantly responding neuron, baseline activity segments were sampled to generate pseudo-trial arrays, repeated 1,000 times. Neurons retained significance if they had significant responses in < 5% of pseudo-trials. **(B)** Trial-averaged normalized response time courses for all significantly responding neurons in each area, as defined in (A), sorted by peak time (note that the same neuron may occur more than once if it responds to more than one photostimulated site). Boxes with dashed outlines indicate the peri-stimulus period when the PMTs were shuttered. **(C)** Average photostimulation-triggered running speeds z scored to pre-stimulation baseline, for MOs and VISp experiments (MOs: n = 140 sites and 5 mice, VISp: n = 190 and 5 mice). Shaded areas: ± S.E.M. **(D)** Distribution of cross-validated linear correlation coefficients (*r*) between halves of the data for MOs neurons using data collected with 5 repeats per stimulated site over 1,000 iterations, compared against distributions from pseudo-trial shuffles. Arrowheads: medians. Printed p-value is from a permutation test. **(E)** Same as (D), but for VISp neurons.

**Figure S4.**
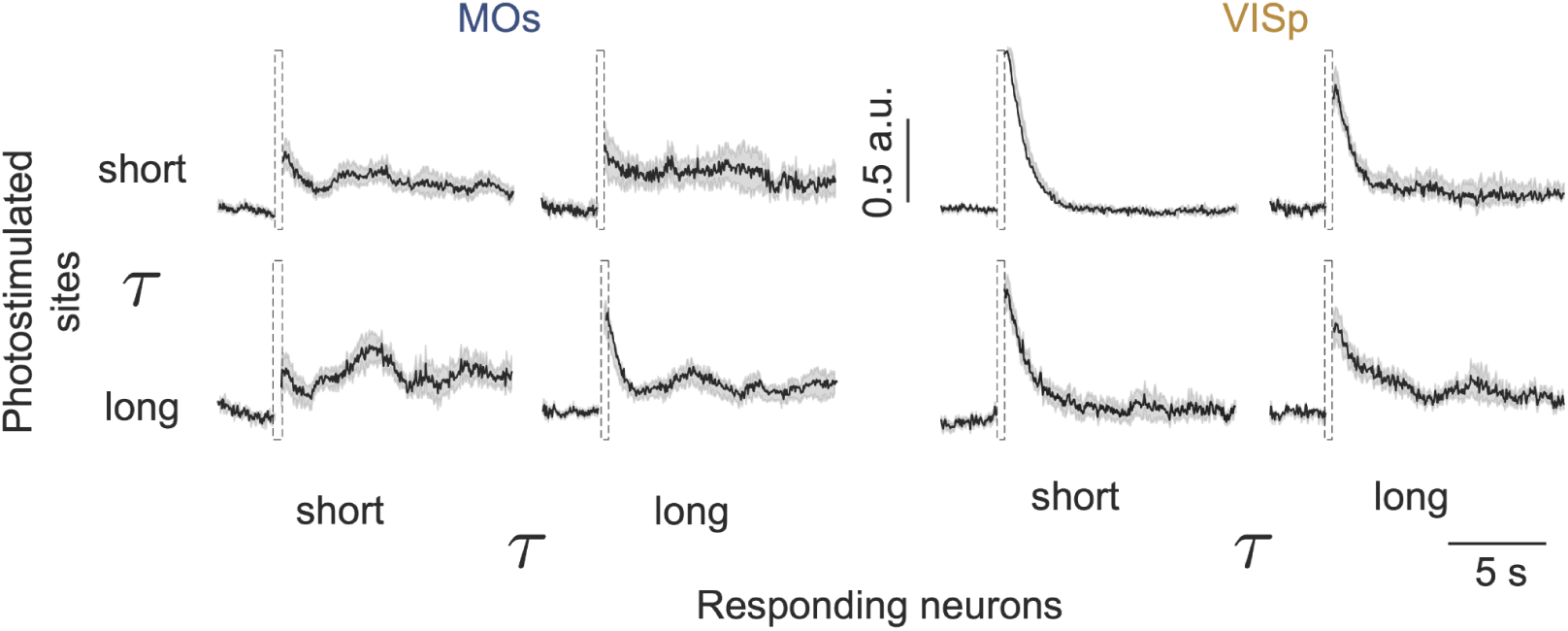
Average population-response timecourses split by timescales of stimulated sites and responding neurons. Peak-normalized photostimulation-triggered responses averaged over all significantly responding neurons in MOs and VISp that also passed inclusion criteria for timescale estimation (i.e., the averages over the rows of the heatmaps in **Fig. 6C**, n = 100neurons and 70 neurons for MOs and VISp, respectively). The data are plotted separately for the different timescale-pair categories defined by median splits. Shaded areas: ± S.E.M.

**Table S1.**
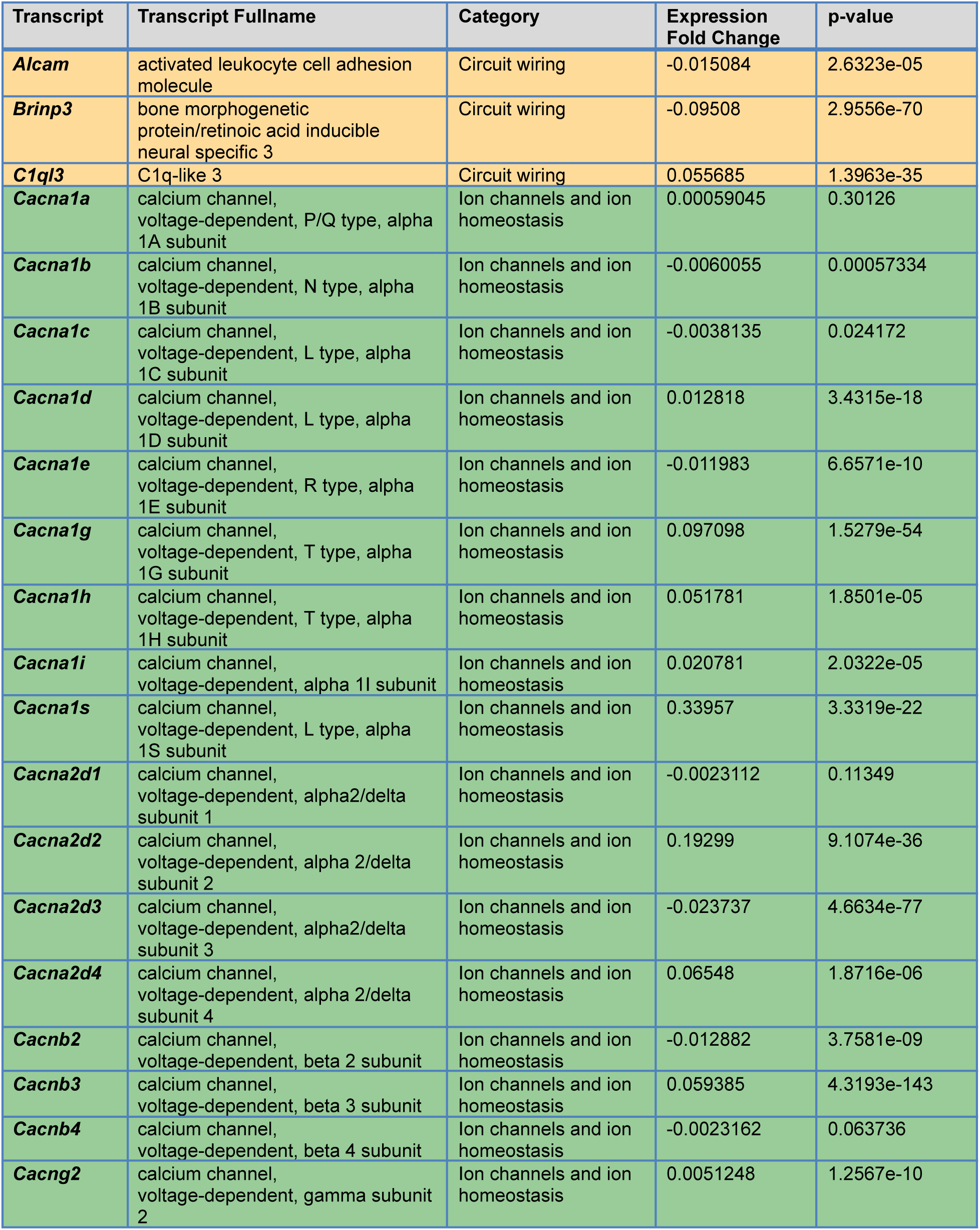
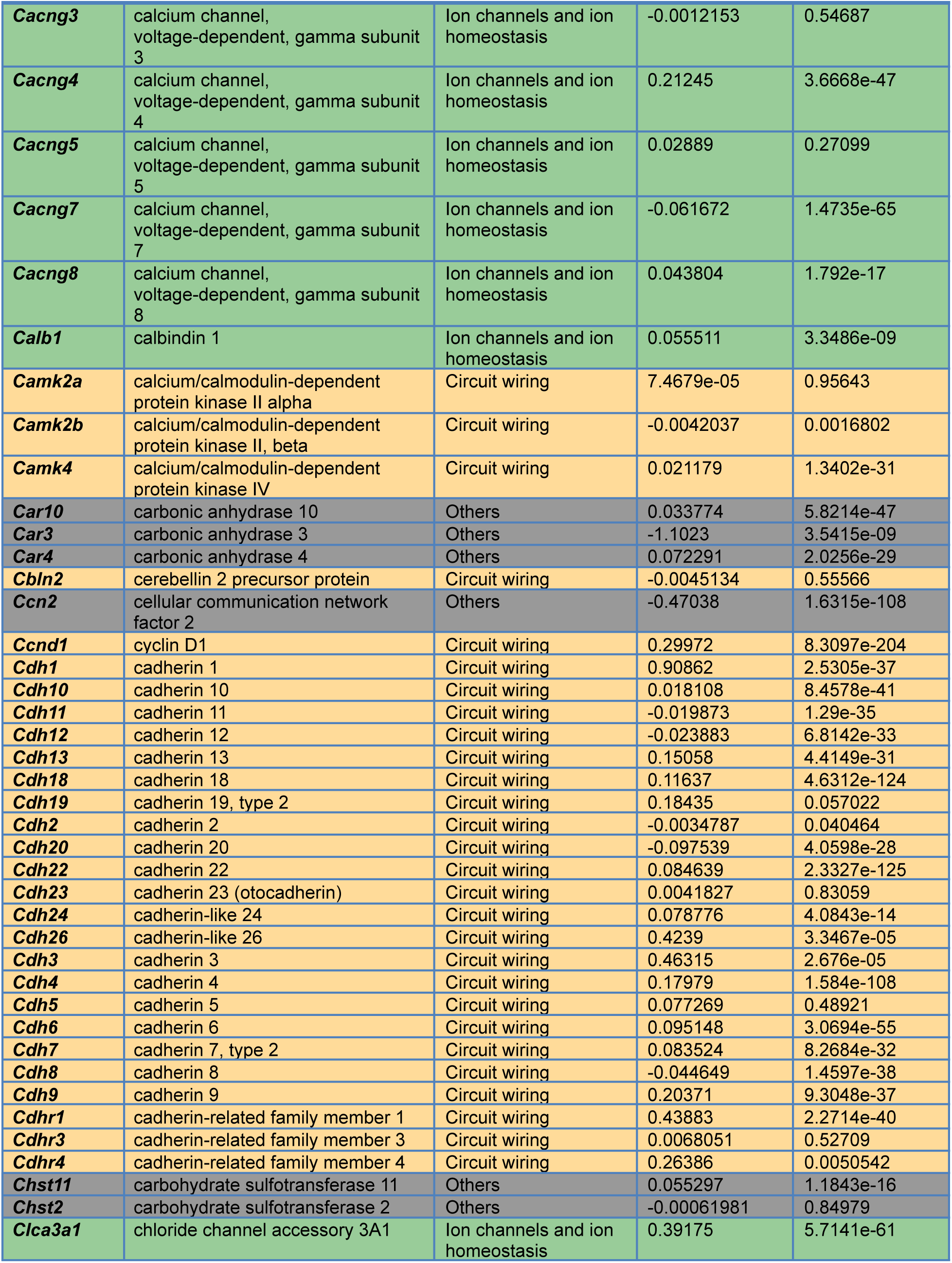
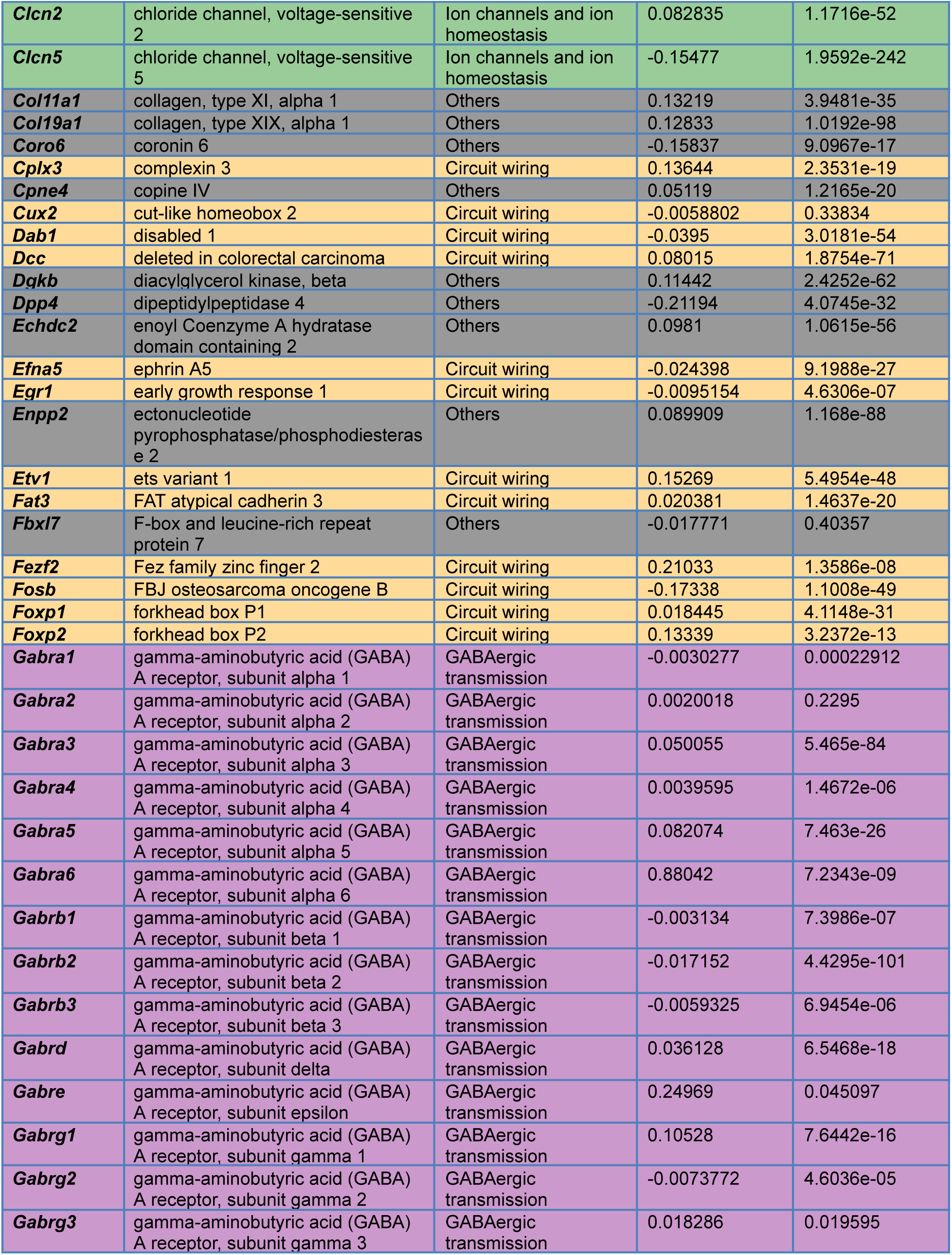
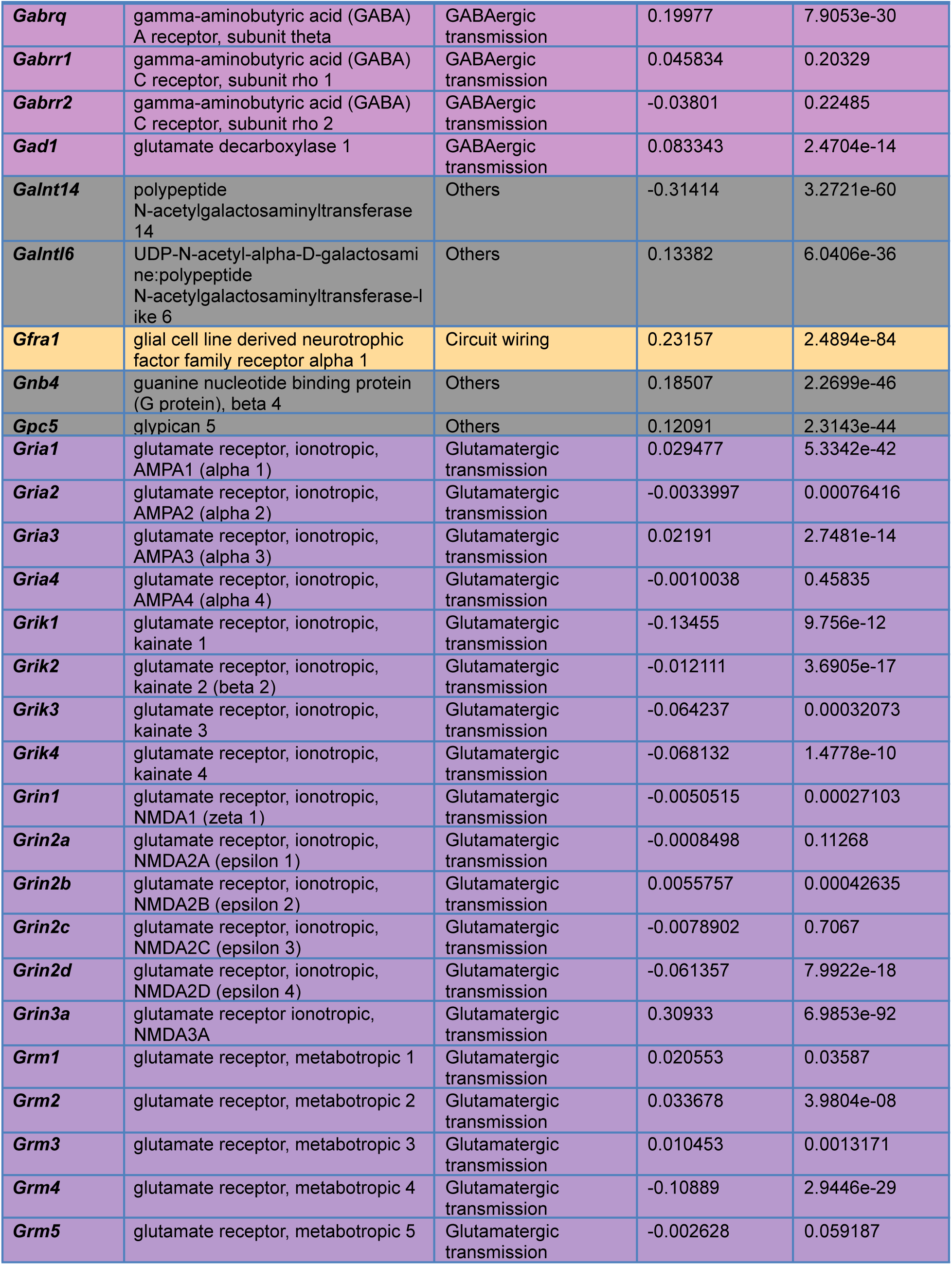
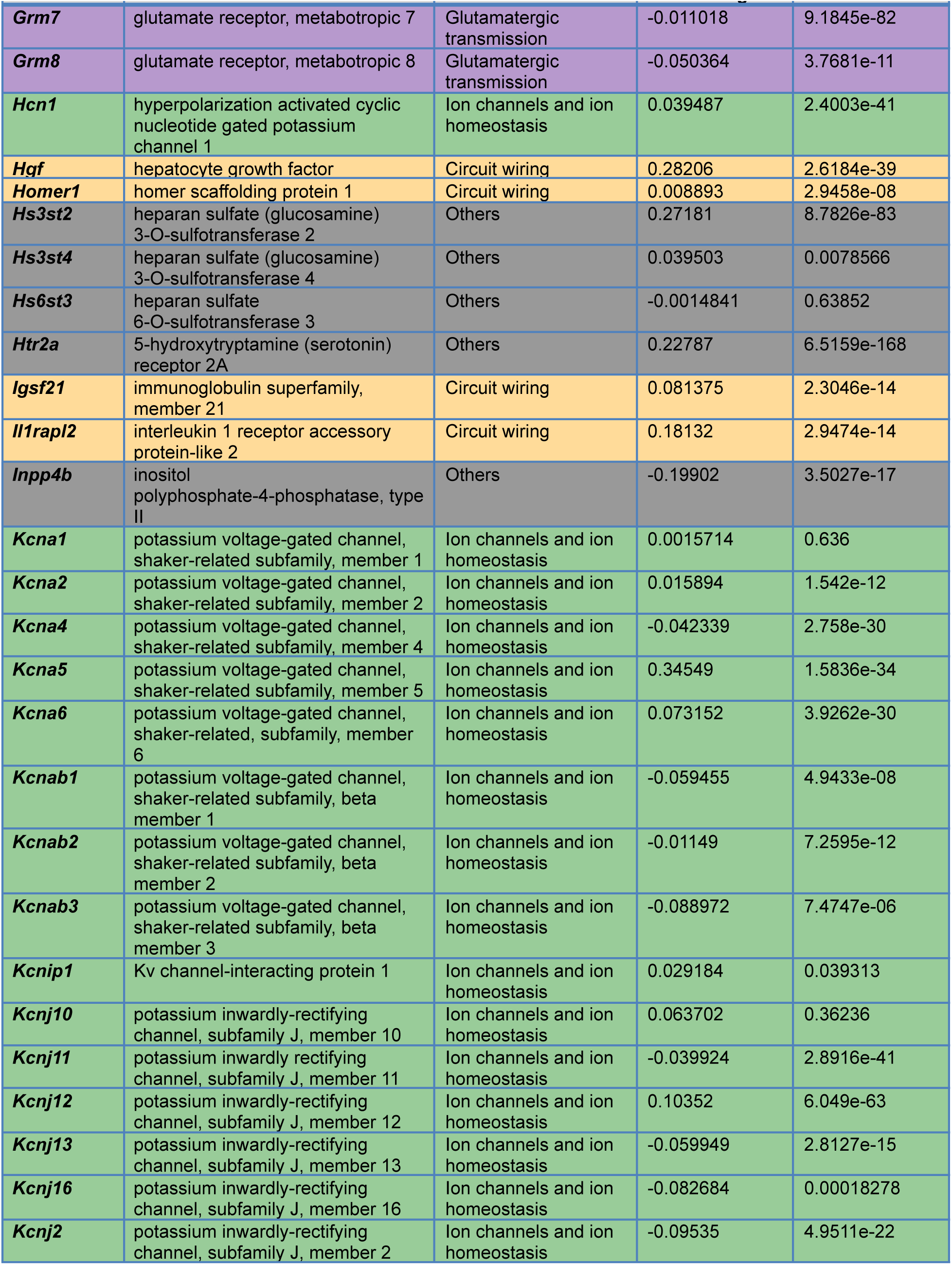
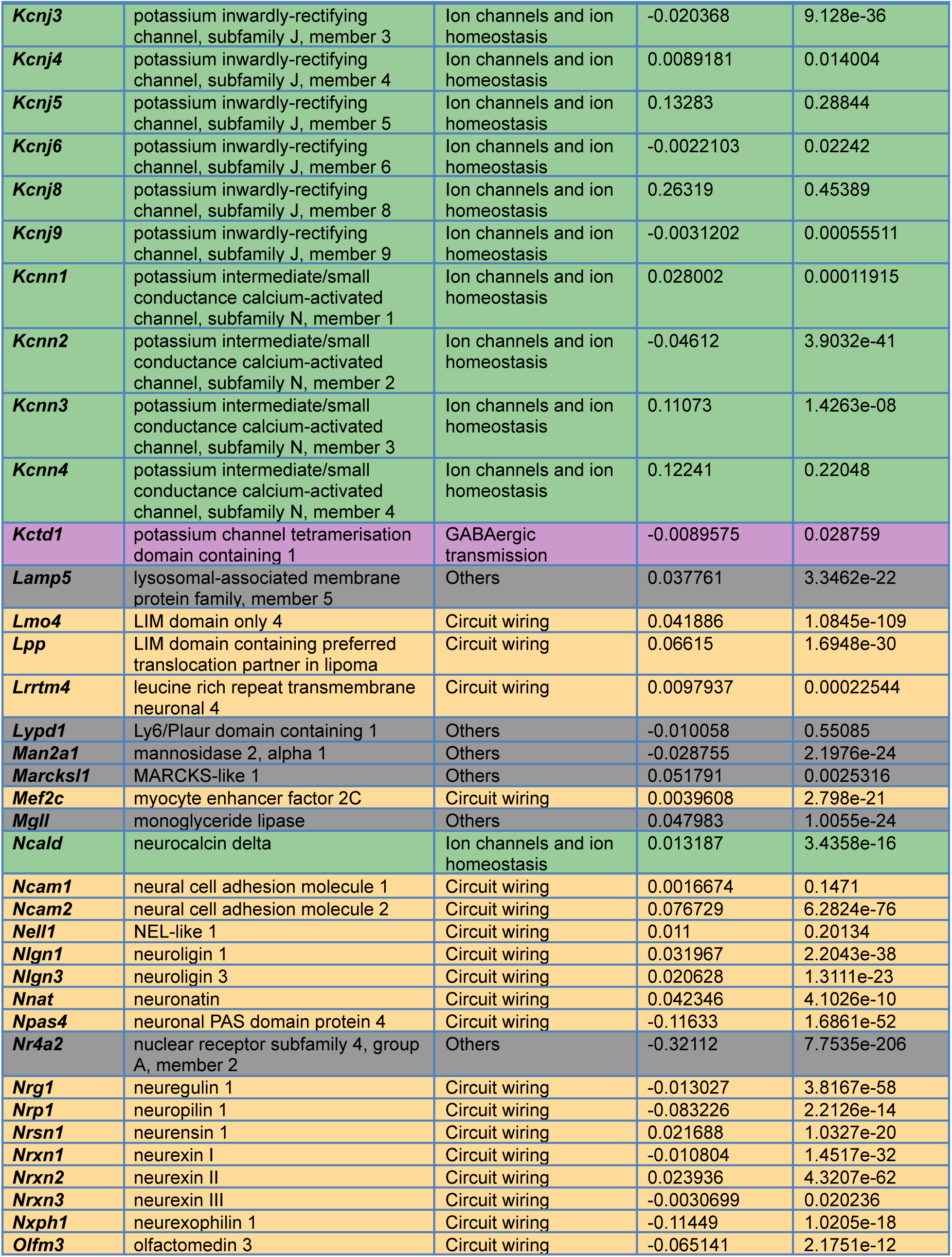
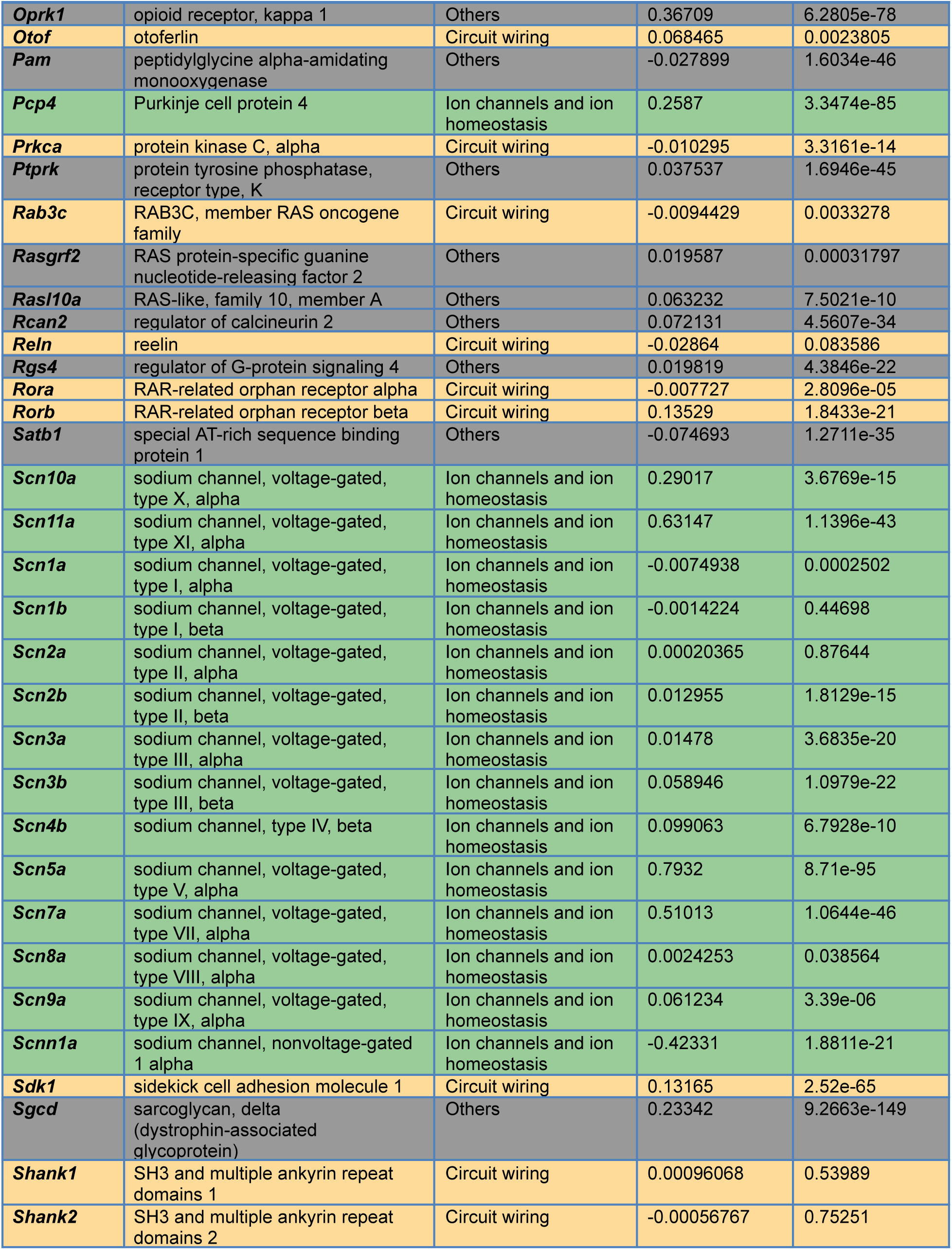
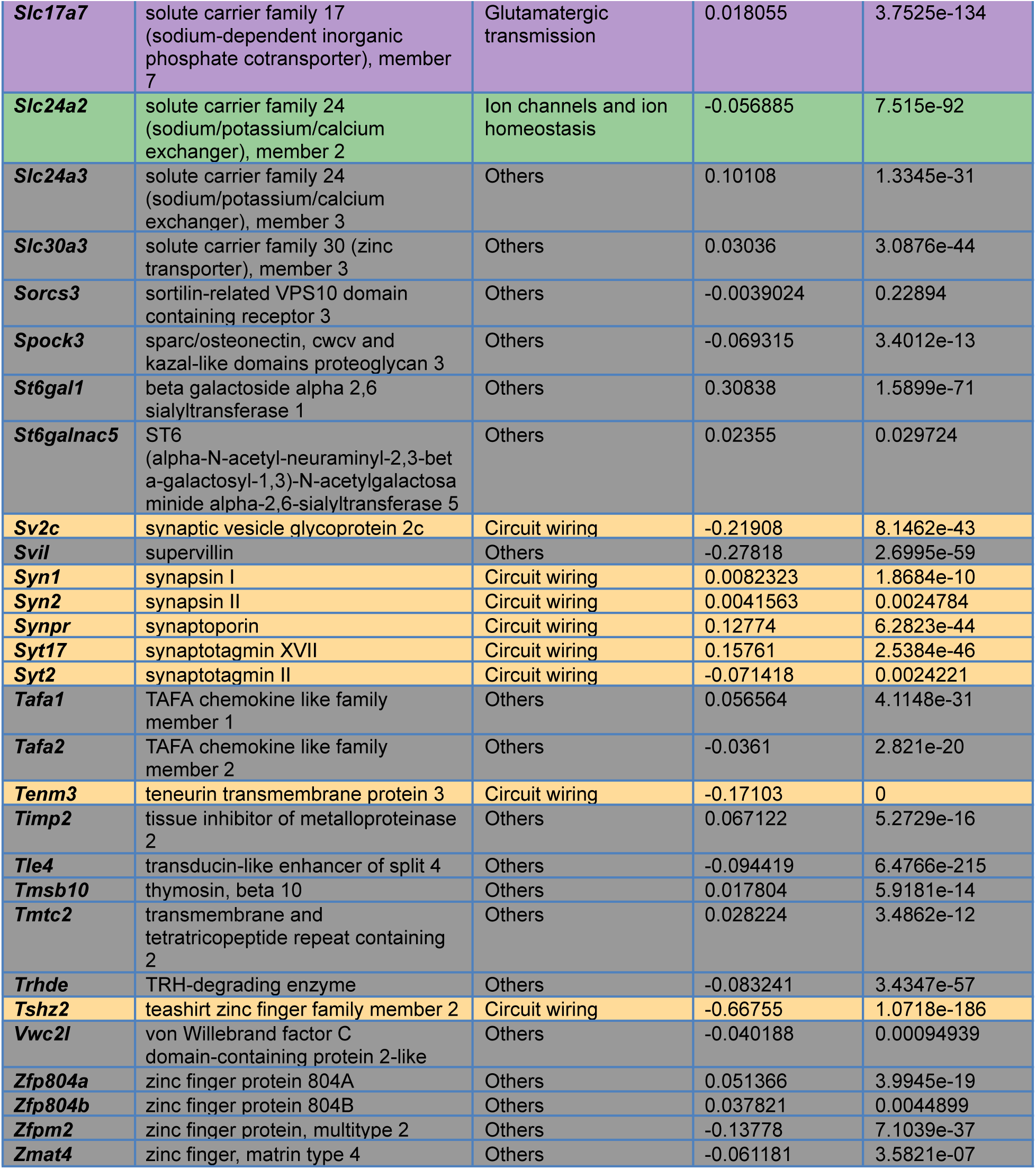
Tabulated data for transcript enrichment analysis in Fig. 1C. Row colors follow transcript category conventions in Figs. 1 and S1. Transcripts are sorted alphabetically.

**Table S2.**
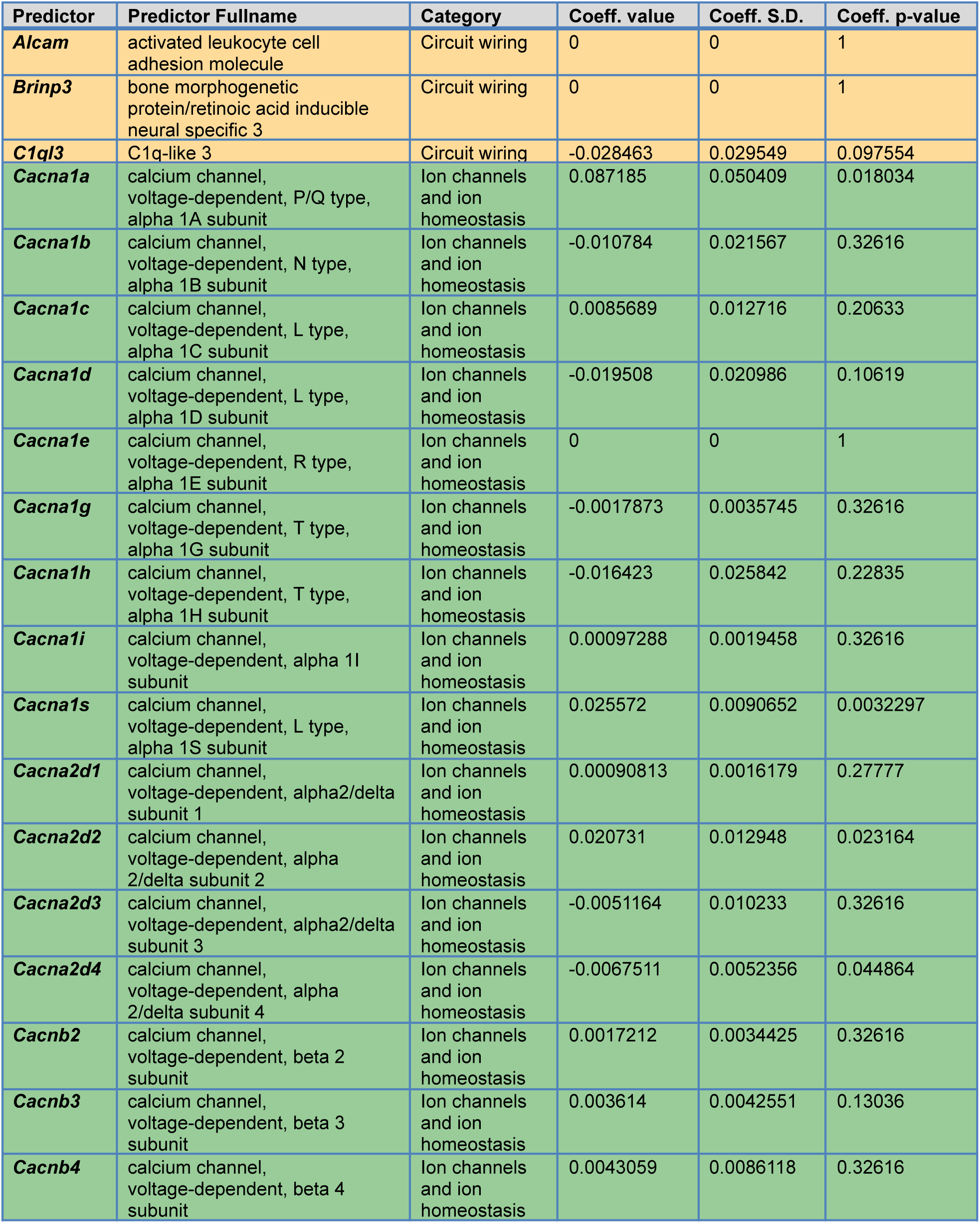
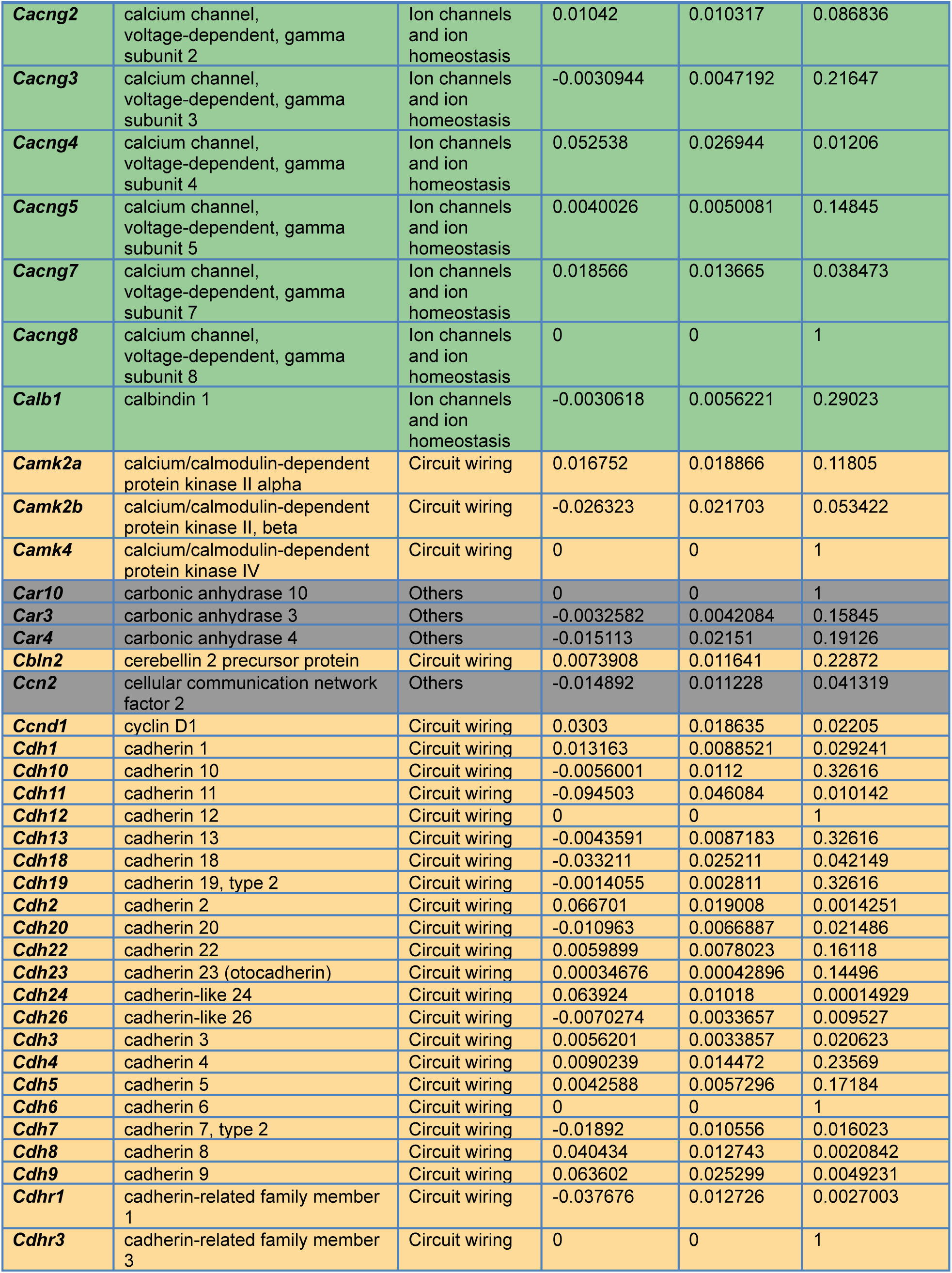
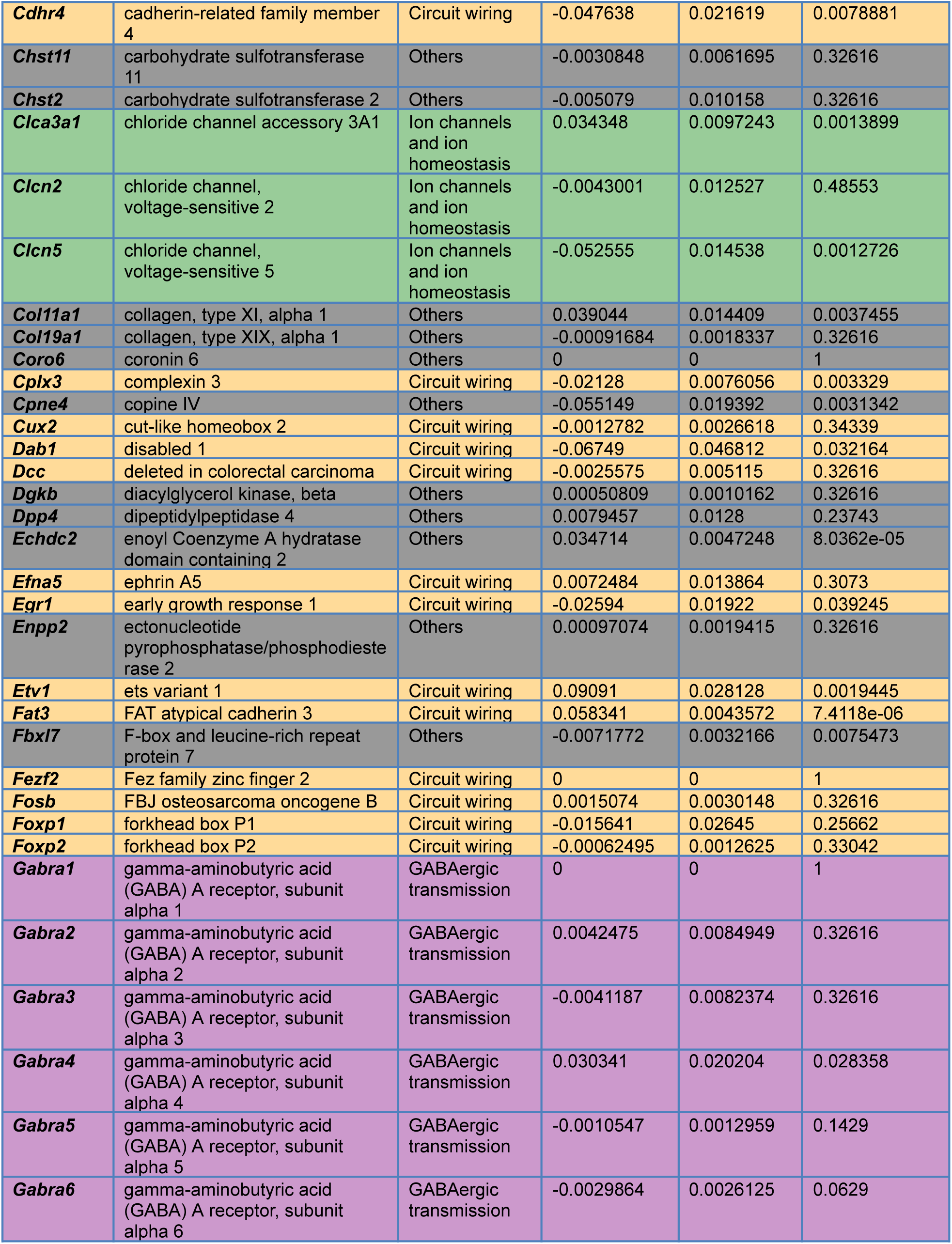
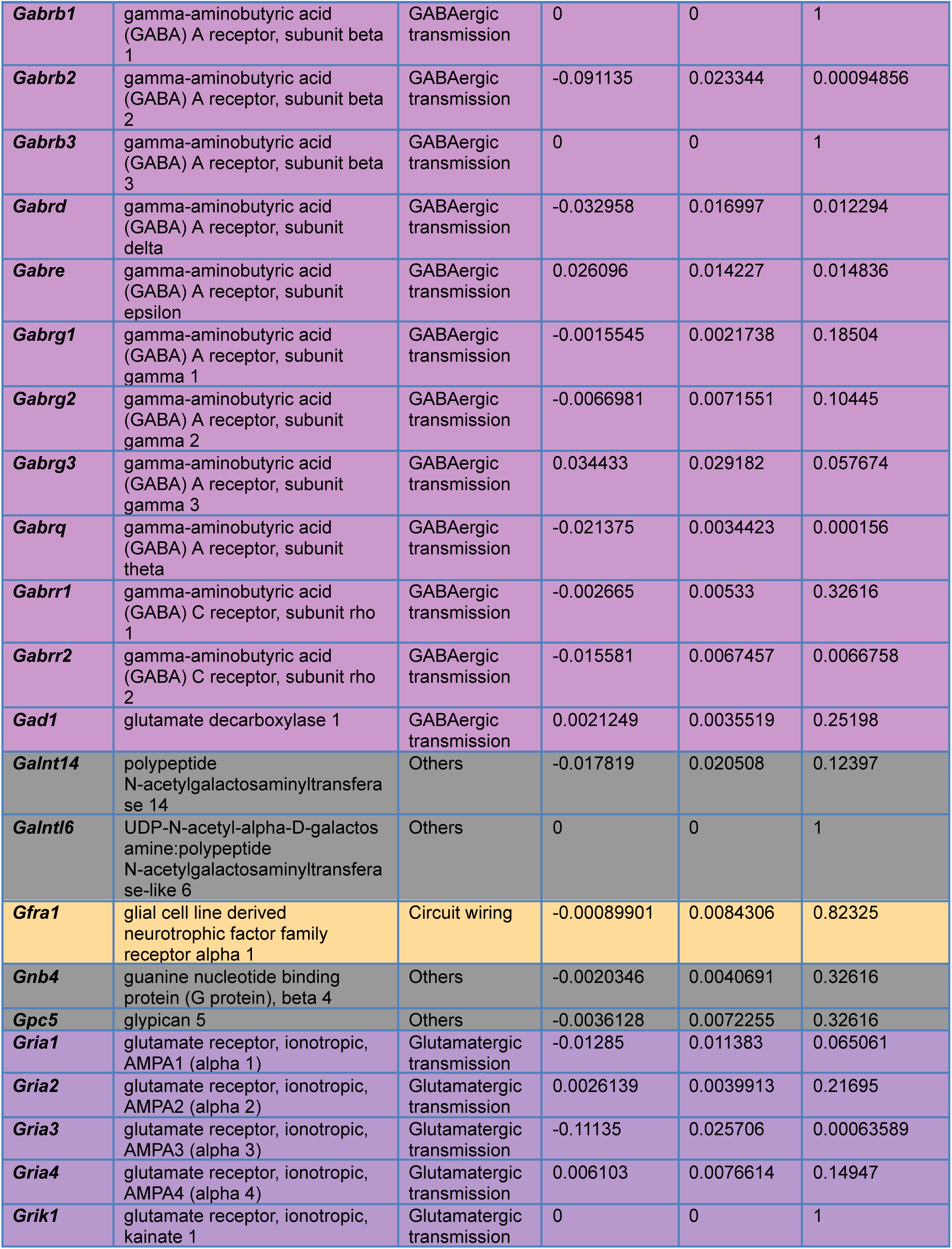
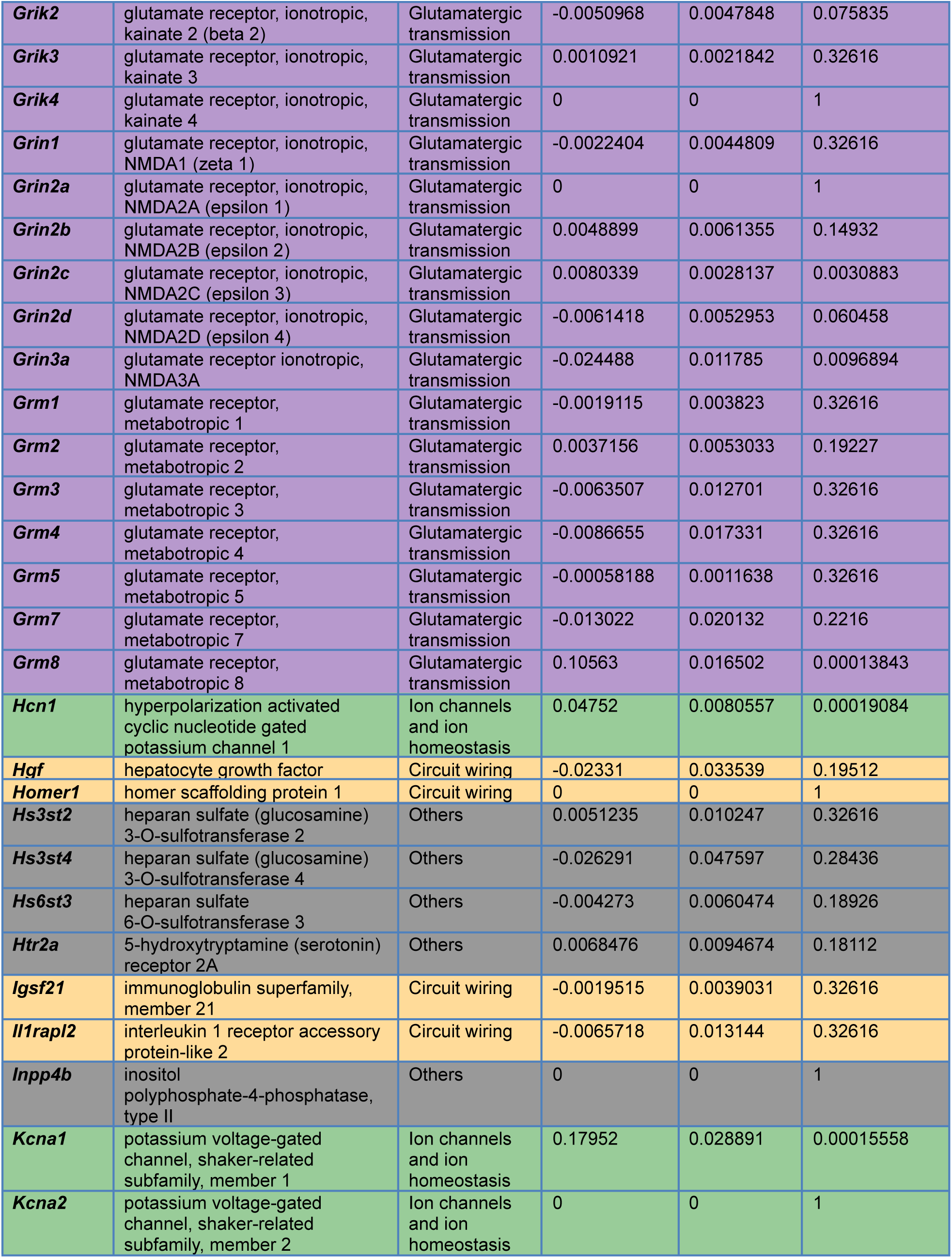
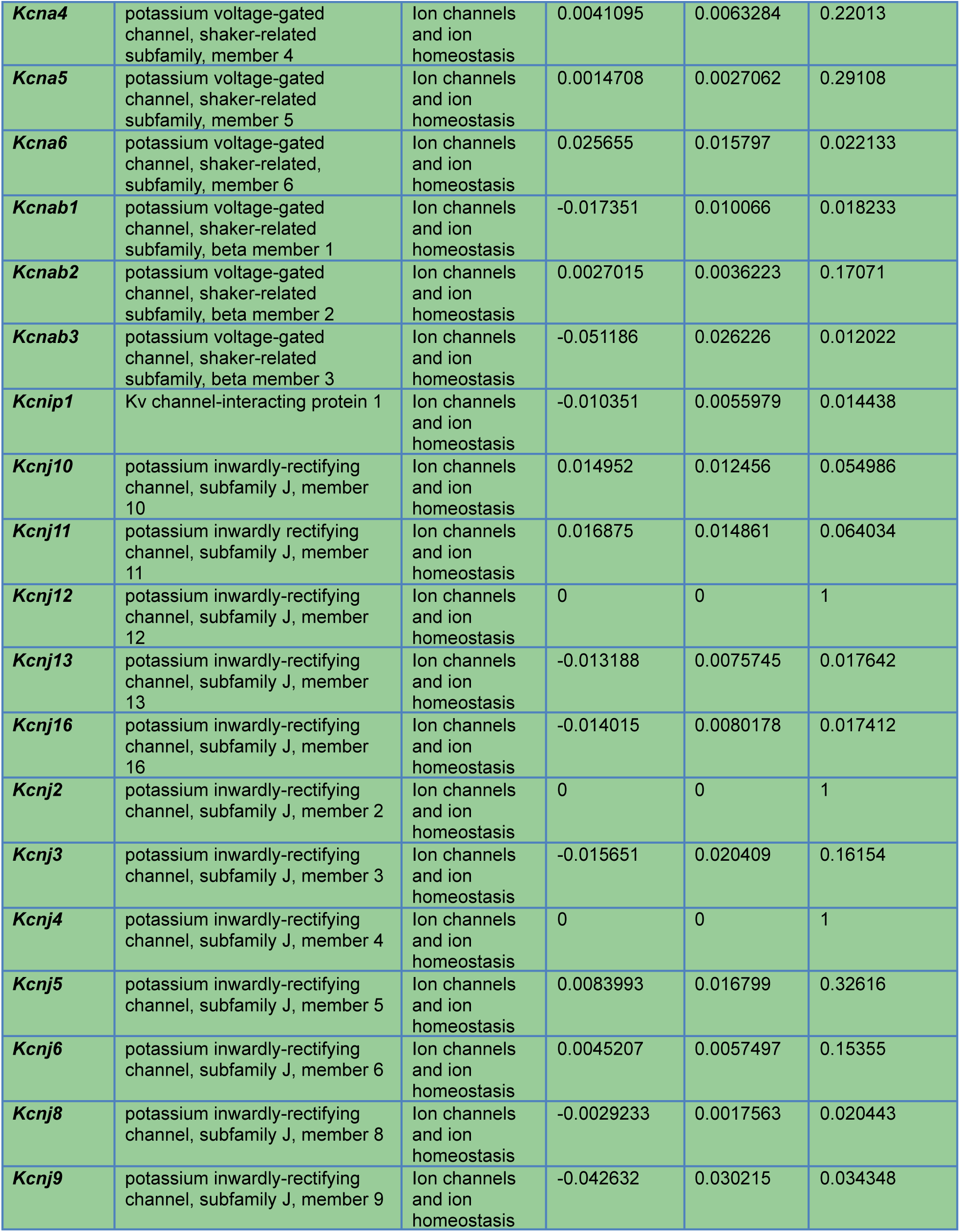
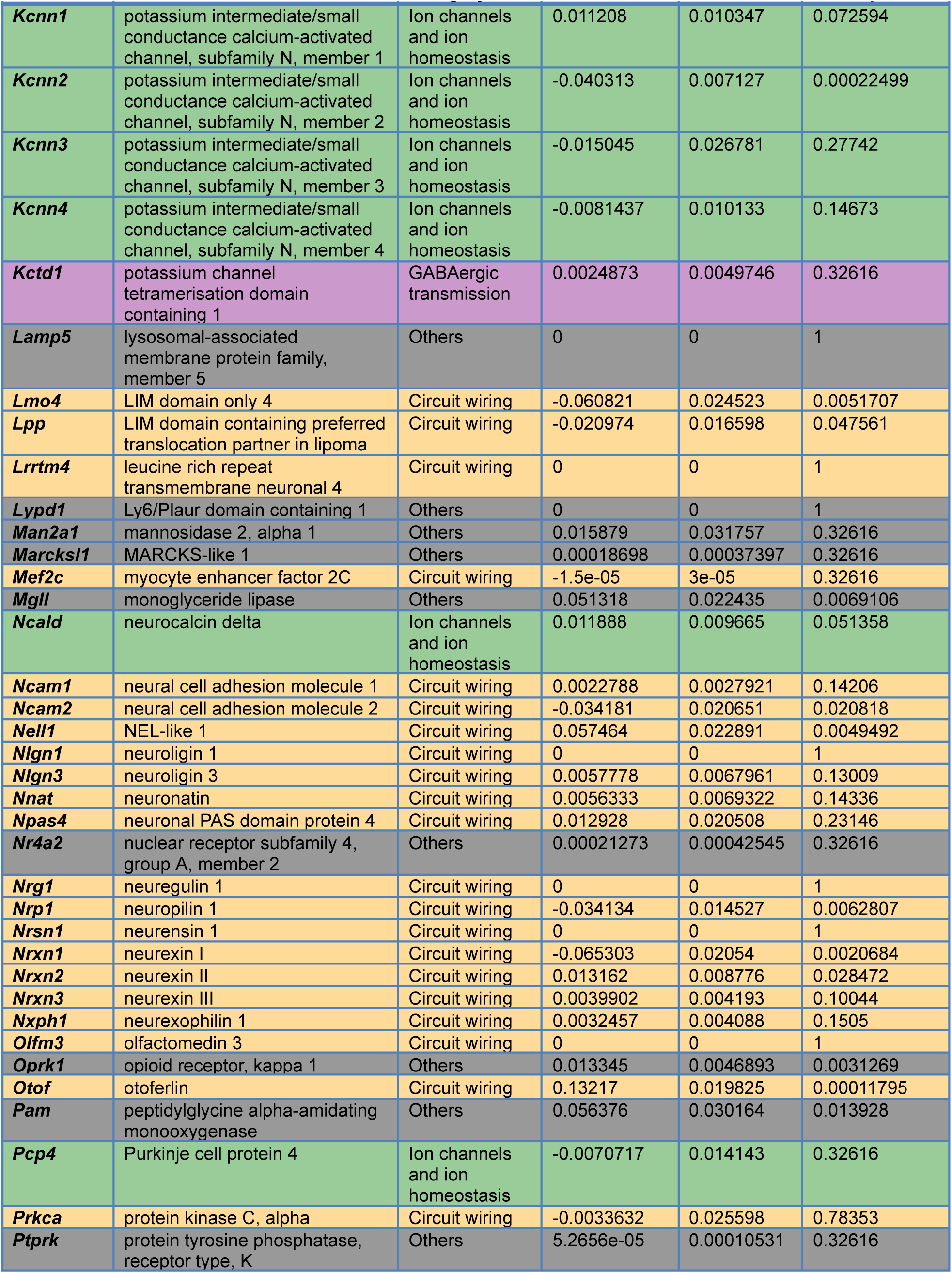
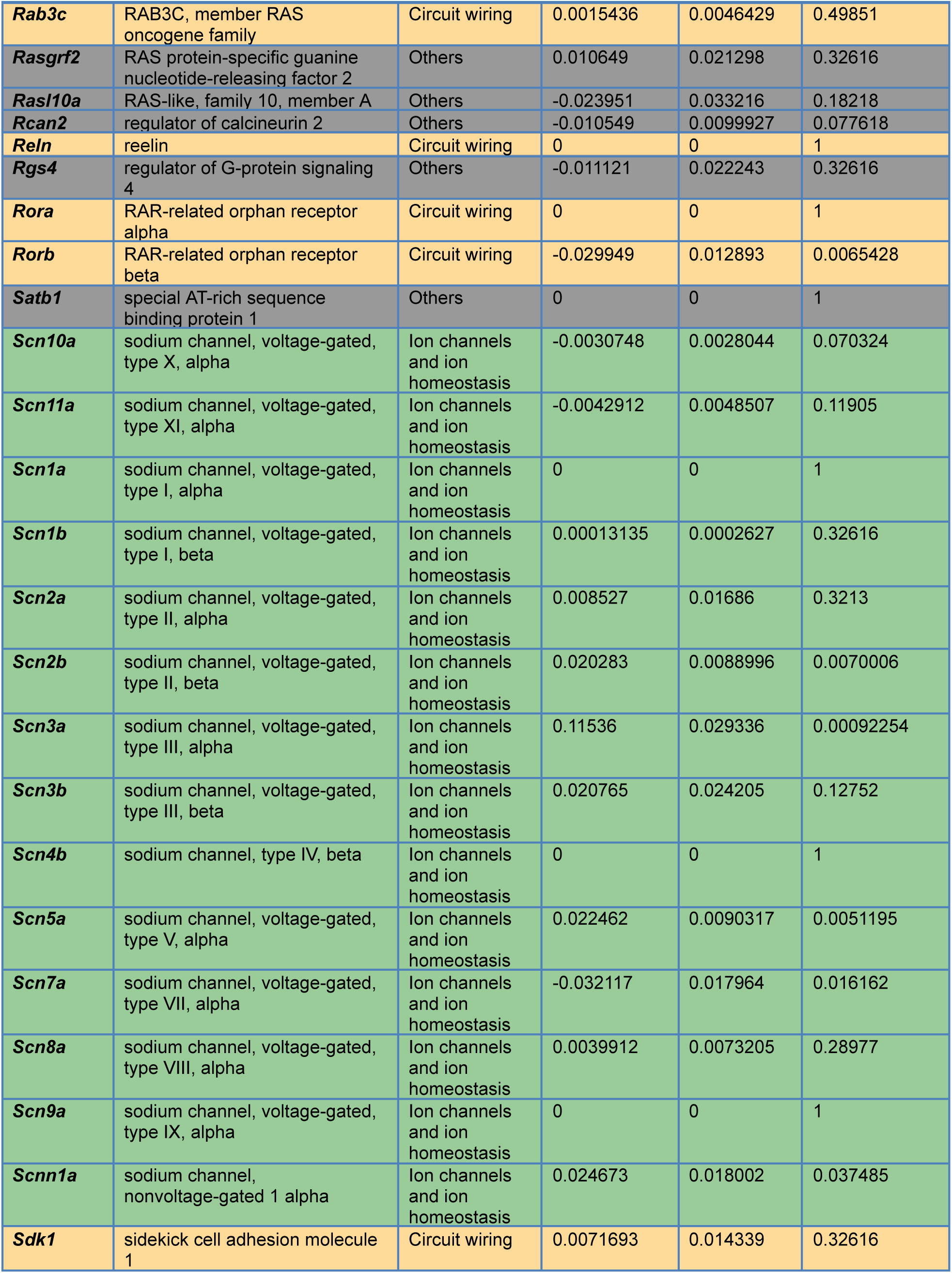
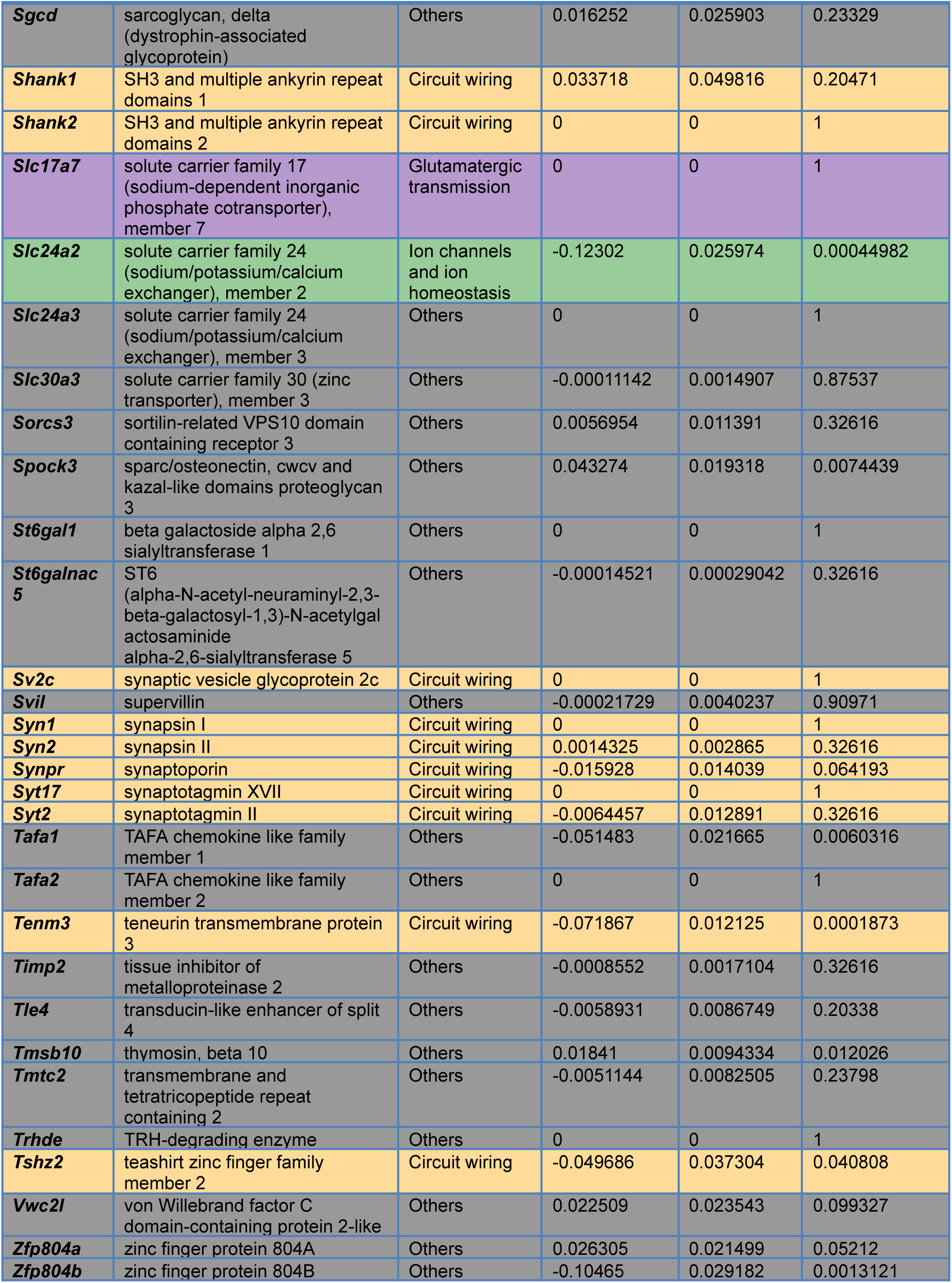
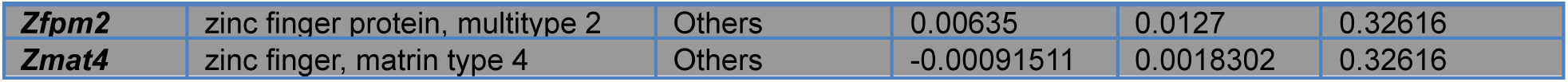
Tabulated data for transcript regression analysis in Fig. 1E. Row colors follow transcript category conventions in Fig. 1. Transcripts are sorted alphabetically.

## Notes

### Competing Interest Statement

The authors have declared no competing interest.

## References

1. Kiebel, S.J., Daunizeau, J., and Friston, K.J. (2008). A hierarchy of time-scales and the brain. PLoS Comput. Biol. 4, e1000209.

2. Murray, J.D., Bernacchia, A., Freedman, D.J., Romo, R., Wallis, J.D., Cai, X., Padoa-Schioppa, C., Pasternak, T., Seo, H., Lee, D., et al. (2014). A hierarchy of intrinsic timescales across primate cortex. Nat. Neurosci. 17, 1661–1663.

3. Gao, R., van den Brink, R.L., Pfeffer, T., and Voytek, B. (2020). Neuronal timescales are functionally dynamic and shaped by cortical microarchitecture. eLife 9, e61277.

4. Manea, A.M.G., Zilverstand, A., Ugurbil, K., Heilbronner, S.R., and Zimmermann, J. (2022). Intrinsic timescales as an organizational principle of neural processing across the whole rhesus macaque brain. eLife 11, e75540.

5. Cavanagh, S.E., Wallis, J.D., Kennerley, S.W., and Hunt, L.T. (2016). Autocorrelation structure at rest predicts value correlates of single neurons during reward-guided choice. eLife 5, e18937.

6. Runyan, C.A., Piasini, E., Panzeri, S., and Harvey, C.D. (2017). Distinct timescales of population coding across cortex. Nature 548, 92–96.

7. Wasmuht, D.F., Spaak, E., Buschman, T.J., Miller, E.K., and Stokes, M.G. (2018). Intrinsic neuronal dynamics predict distinct functional roles during working memory. Nat. Commun. 9, 3499.

8. Cavanagh, S.E., Towers, J.P., Wallis, J.D., Hunt, L.T., and Kennerley, S.W. (2018). Reconciling persistent and dynamic hypotheses of working memory coding in prefrontal cortex. Nat. Commun. 9, 3498.

9. Ito, T., Hearne, L.J., and Cole, M.W. (2020). A cortical hierarchy of localized and distributed processes revealed via dissociation of task activations, connectivity changes, and intrinsic timescales. NeuroImage 221, 117141.

10. Soltani, A., Murray, J.D., Seo, H., and Lee, D. (2021). Timescales of cognition in the brain. Curr. Opin. Behav. Sci. 41, 30–37.

11. Murphy, P.R., Wilming, N., Hernandez-Bocanegra, D.C., Prat-Ortega, G., and Donner, T.H. (2021). Adaptive circuit dynamics across human cortex during evidence accumulation in changing environments. Nat. Neurosci. 24, 987–997.

12. Siegle, J.H., Jia, X., Durand, S., Gale, S., Bennett, C., Graddis, N., Heller, G., Ramirez, T.K., Choi, H., Luviano, J.A., et al. (2021). Survey of spiking in the mouse visual system reveals functional hierarchy. Nature 592, 86–92.

13. Zeraati, R., Shi, Y.-L., Steinmetz, N.A., Gieselmann, M.A., Thiele, A., Moore, T., Levina, A., and Engel, T.A. (2023). Intrinsic timescales in the visual cortex change with selective attention and reflect spatial connectivity. Nat. Commun. 14, 1858.

14. Costa, R.M., Luo, J.K., Salvino, P.S., Ackert-Smith, L.A., Ibarra, S.M., and Pinto, L. (2025). Cognitive processes are disentangled at cortex-wide scales. bioRxiv, 2025.07.24.666672. 10.1101/2025.07.24.666672.

15. Hasson, U., Yang, E., Vallines, I., Heeger, D.J., and Rubin, N. (2008). A hierarchy of temporal receptive windows in human cortex. J. Neurosci. 28, 2539–2550.

16. Orsolic, I., Rio, M., Mrsic-Flogel, T.D., and Znamenskiy, P. (2021). Mesoscale cortical dynamics reflect the interaction of sensory evidence and temporal expectation during perceptual decision-making. Neuron 109, 1861–1875.e10.

17. Pinto, L., Tank, D.W., and Brody, C.D. (2022). Multiple timescales of sensory-evidence accumulation across the dorsal cortex. eLife 11, e70263.

18. Wang, X.-J. (2022). Theory of the Multiregional Neocortex: Large-Scale Neural Dynamics and Distributed Cognition. Annu. Rev. Neurosci. 45, 533–560.

19. Ballesteros-Yáñez, I., Benavides-Piccione, R., Elston, G.N., Yuste, R., and DeFelipe, J. (2006). Density and morphology of dendritic spines in mouse neocortex. Neuroscience 138, 403–409.

20. Burt, J.B., Demirtaş, M., Eckner, W.J., Navejar, N.M., Ji, J.L., Martin, W.J., Bernacchia, A., Anticevic, A., and Murray, J.D. (2018). Hierarchy of transcriptomic specialization across human cortex captured by structural neuroimaging topography. Nat. Neurosci. 21, 1251–1259.

21. Fulcher, B.D., Murray, J.D., Zerbi, V., and Wang, X.-J. (2019). Multimodal gradients across mouse cortex. Proc. Natl. Acad. Sci. U. S. A. 116, 4689–4695.

22. Wang, X.-J. (2020). Macroscopic gradients of synaptic excitation and inhibition in the neocortex. Nat. Rev. Neurosci. 21, 169–178.

23. Duarte, R., Seeholzer, A., Zilles, K., and Morrison, A. (2017). Synaptic patterning and the timescales of cortical dynamics. Curr. Opin. Neurobiol. 43, 156–165.

24. Bernacchia, A., Seo, H., Lee, D., and Wang, X.-J. (2011). A reservoir of time constants for memory traces in cortical neurons. Nat. Neurosci. 14, 366–372.

25. Zylberberg, J., and Strowbridge, B.W. (2017). Mechanisms of persistent activity in cortical circuits: Possible neural substrates for working memory. Annu. Rev. Neurosci. 40, 603–627.

26. Zeraati, R., Levina, A., Macke, J.H., and Gao, R. (2024). Neural timescales from a computational perspective. arXiv:2409.02684.

27. Khajehabdollahi, S., Zeraati, R., Giannakakis, E., Schäfer, T.J., Martius, G., and Levina, A. (2023). Emergent mechanisms for long timescales depend on training curriculum and affect performance in memory tasks. arXiv:2309.12927

28. Chaudhuri, R., Knoblauch, K., Gariel, M.-A., Kennedy, H., and Wang, X.-J. (2015). A Large-Scale Circuit Mechanism for Hierarchical Dynamical Processing in the Primate Cortex. Neuron 88, 419–431.

29. Deco, G., Kringelbach, M.L., Arnatkeviciute, A., Oldham, S., Sabaroedin, K., Rogasch, N.C., Aquino, K.M., and Fornito, A. (2021). Dynamical consequences of regional heterogeneity in the brain’s transcriptional landscape. Sci. Adv. 7, eabf4752.

30. Mejías, J.F., and Wang, X.-J. (2022). Mechanisms of distributed working memory in a large-scale network of macaque neocortex. eLife 11, e72136.

31. Mante, V., Sussillo, D., Shenoy, K.V., and Newsome, W.T. (2013). Context-dependent computation by recurrent dynamics in prefrontal cortex. Nature 503, 78–84.

32. Reinhold, K., Lien, A.D., and Scanziani, M. (2015). Distinct recurrent versus afferent dynamics in cortical visual processing. Nat. Neurosci. 18, 1789–1797.

33. Inagaki, H.K., Fontolan, L., Romani, S., and Svoboda, K. (2019). Discrete attractor dynamics underlies persistent activity in the frontal cortex. Nature 566, 212–217.

34. Peron, S., Pancholi, R., Voelcker, B., Wittenbach, J.D., Ólafsdóttir, H.F., Freeman, J., and Svoboda, K. (2020). Recurrent interactions in local cortical circuits. Nature 579, 256–259.

35. Daie, K., Svoboda, K., and Druckmann, S. (2021). Targeted photostimulation uncovers circuit motifs supporting short-term memory. Nat. Neurosci. 24, 259–265.

36. O’Rawe, J.F., Zhou, Z., Li, A.J., LaFosse, P.K., Goldbach, H.C., and Histed, M.H. (2023). Excitation creates a distributed pattern of cortical suppression due to varied recurrent input. Neuron 111, 4086–4101.

37. Oldenburg, I.A., Hendricks, W.D., Handy, G., Shamardani, K., Bounds, H.A., Doiron, B., and Adesnik, H. (2024). The logic of recurrent circuits in the primary visual cortex. Nat. Neurosci. 27, 137–147.

38. Zhang, M., Pan, X., Jung, W., Halpern, A.R., Eichhorn, S.W., Lei, Z., Cohen, L., Smith, K.A., Tasic, B., Yao, Z., et al. (2023). Molecularly defined and spatially resolved cell atlas of the whole mouse brain. Nature 624, 343–354.

39. Chen, X., Fischer, S., Rue, M.C.P., Zhang, A., Mukherjee, D., Kanold, P.O., Gillis, J., and Zador, A.M. (2024). Whole-cortex in situ sequencing reveals input-dependent area identity. Nature, 10.1038/s41586-024-07221-6.

40. Arikkath, J., and Reichardt, L.F. (2008). Cadherins and catenins at synapses: roles in synaptogenesis and synaptic plasticity. Trends Neurosci. 31, 487–494.

41. Sompolinsky, H., Crisanti, A., and Sommers, H.J. (1988). Chaos in random neural networks. Phys. Rev. Lett. 61, 259–262.

42. Barthas, F., and Kwan, A.C. (2017). Secondary motor cortex: where “sensory” meets “motor” in the rodent frontal cortex. Trends Neurosci. 40, 181–193.

43. Khilkevich, A., Lohse, M., Low, R., Orsolic, I., Bozic, T., Windmill, P., and Mrsic-Flogel, T.D. (2024). Brain-wide dynamics linking sensation to action during decision-making. Nature 634, 890–900 (2024).

44. Imani, E., Radkani, S., Hashemi, A., Harati, A., Pourreza, H., and Moazami Goudarzi, M. (2023). Distributed coding of evidence accumulation across the mouse brain using microcircuits with a diversity of timescales. eNeuro 10, ENEURO.0282–23.2023.

45. Dombeck, D.A., Graziano, M.S., and Tank, D.W. (2009). Functional Clustering of Neurons in Motor Cortex Determined by Cellular Resolution Imaging in Awake Behaving Mice. J. Neurosci. 29, 13751–13760.

46. Pinto, L., and Dan, Y. (2015). Cell-Type-Specific Activity in Prefrontal Cortex during Goal-Directed Behavior. Neuron 87, 437–450.

47. Ringach, D.L., Mineault, P.J., Tring, E., Olivas, N.D., Garcia-Junco-Clemente, P., and Trachtenberg, J.T. (2016). Spatial clustering of tuning in mouse primary visual cortex. Nat. Commun. 7, 12270.

48. Gu, Y., Lewallen, S., Kinkhabwala, A.A., Domnisoru, C., Yoon, K., Gauthier, J.L., Fiete, I.R., and Tank, D.W. (2018). A map-like micro-organization of grid cells in the medial entorhinal cortex. Cell 175, 736–750.e30.

49. Chang, H., Esteves, I.M., Neumann, A.R., Mohajerani, M.H., and McNaughton, B.L. (2023). Cortical reactivation of spatial and non-spatial features coordinates with hippocampus to form a memory dialogue. Nat. Commun. 14, 7748.

50. Issa, J.B., Haeffele, B.D., Agarwal, A., Bergles, D.E., Young, E.D., and Yue, D.T. (2014). Multiscale Optical Ca2+ Imaging of Tonal Organization in Mouse Auditory Cortex. Neuron 83, 944–959.

51. Rickgauer, J.P., Deisseroth, K., and Tank, D.W. (2014). Simultaneous cellular-resolution optical perturbation and imaging of place cell firing fields. Nat. Neurosci. 17, 1816–1824.

52. Packer, A.M., Russell, L.E., Dalgleish, H.W.P., and Hausser, M.A. (2015). Simultaneous all-optical manipulation and recording of neural circuit activity with cellular resolution in vivo. Nat. Methods 12, 140–146.

53. Chettih, S.N., and Harvey, C.D. (2019). Single-neuron perturbations reveal feature-specific competition in V1. Nature 567, 334–340.

54. Lees, R.M., Pichler, B., and Packer, A.M. (2023). Optical resolution is not the limiting factor for spatial precision of two-photon optogenetic photostimulation. bioRxiv, 2023.07.01.547318. 10.1101/2023.07.01.547318.

55. Herculano-Houzel, S., Watson, C., and Paxinos, G. (2013). Distribution of neurons in functional areas of the mouse cerebral cortex reveals quantitatively different cortical zones. Front. Neuroanat. 7, 35.

56. Luczak, A., Barthó, P., Marguet, S.L., Buzsáki, G., and Harris, K.D. (2007). Sequential structure of neocortical spontaneous activity in vivo. Proc. Natl. Acad. Sci. U. S. A. 104, 347–352.

57. Harvey, C.D., Coen, P., and Tank, D.W. (2012). Choice-specific sequences in parietal cortex during a virtual-navigation decision task. Nature 484, 62–68.

58. Akhlaghpour, H., Wiskerke, J., Choi, J.Y., Taliaferro, J.P., Au, J., and Witten, I.B. (2016). Dissociated sequential activity and stimulus encoding in the dorsomedial striatum during spatial working memory. eLife 5, e19507.

59. Koay, S.A., Charles, A.S., Thiberge, S.Y., Brody, C.D., and Tank, D.W. (2022). Sequential and efficient neural-population coding of complex task information. Neuron 110, 328–349.e11.

60. Bigus, E.R., Lee, H.-W., Bowler, J.C., Shi, J., and Heys, J.G. (2024). Medial entorhinal cortex mediates learning of context-dependent interval timing behavior. Nat. Neurosci. 27, 1587–1598.

61. Morcos, A.S., and Harvey, C.D. (2016). History-dependent variability in population dynamics during evidence accumulation in cortex. Nat. Neurosci. 19, 1672–1681.

62. Nieh, E.H., Schottdorf, M., Freeman, N.W., Low, R.J., Lewallen, S., Koay, S.A., Pinto, L., Gauthier, J.L., Brody, C.D., and Tank, D.W. (2021). Geometry of abstract learned knowledge in the hippocampus. Nature 595, 80–84.

63. Buzsáki, G., and Tingley, D. (2018). Space and time: The hippocampus as a sequence generator. Trends Cogn. Sci. 22, 853–869.

64. Enel, P., Wallis, J.D., and Rich, E.L. (2020). Stable and dynamic representations of value in the prefrontal cortex. eLife 9, e54313.

65. Parker, N.F., Baidya, A., Cox, J., Haetzel, L.M., Zhukovskaya, A., Murugan, M., Engelhard, B., Goldman, M.S., and Witten, I.B. (2022). Choice-selective sequences dominate in cortical relative to thalamic inputs to NAc to support reinforcement learning. Cell Rep. 39, 110756.

66. Curto, C., Sakata, S., Marguet, S., Itskov, V., and Harris, K.D. (2009). A simple model of cortical dynamics explains variability and state dependence of sensory responses in urethane-anesthetized auditory cortex. J. Neurosci. 29, 10600–10612.

67. Kaufman, M.T., Churchland, M.M., Ryu, S.I., and Shenoy, K.V. (2014). Cortical activity in the null space: permitting preparation without movement. Nat. Neurosci. 17, 440–448.

68. Wu, A., Chen, Q., Yu, B., Chae, S., Tan, Z., Ramot, A., and Komiyama, T. (2025). Targeted stimulation of motor cortex neural ensembles drives learned movements. bioRxiv 2025.01.06.631504; doi: 10.1101/2025.01.06.631504.

69. Scott, B.B., Constantinople, C.M., Akrami, A., Hanks, T.D., Brody, C.D., and Tank, D.W. (2017). Fronto-parietal Cortical Circuits Encode Accumulated Evidence with a Diversity of Timescales. Neuron 95, 385–398.e5.

70. Spellman, T., Rigotti, M., Ahmari, S.E., Fusi, S., Gogos, J.A., and Gordon, J.A. (2015). Hippocampal–prefrontal input supports spatial encoding in working memory. Nature 522, 309–314.

71. Bolkan, S.S., Stujenske, J.M., Parnaudeau, S., Spellman, T.J., Rauffenbart, C., Abbas, A.I., Harris, A.Z., Gordon, J.A., and Kellendonk, C. (2017). Thalamic projections sustain prefrontal activity during working memory maintenance. Nat. Neurosci. 20, 987–996.

72. Guo, Z.V., Inagaki, H.K., Daie, K., Druckmann, S., Gerfen, C.R., and Svoboda, K. (2017). Maintenance of persistent activity in a frontal thalamocortical loop. Nature 545, 181–186.

73. Voitov, I., and Mrsic-Flogel, T.D. (2022). Cortical feedback loops bind distributed representations of working memory. Nature 608, 381–389.

74. Kepecs, A., and Fishell, G. (2014). Interneuron cell types are fit to function. Nature 505, 318–326.

75. Wang, Y., Markram, H., Goodman, P.H., Berger, T.K., Ma, J., and Goldman-Rakic, P.S. (2006). Heterogeneity in the pyramidal network of the medial prefrontal cortex. Nat. Neurosci. 9, 534–542.

76. Varela, J.A., Sen, K., Gibson, J., Fost, J., Abbott, L.F., and Nelson, S.B. (1997). A quantitative description of short-term plasticity at excitatory synapses in layer 2/3 of rat primary visual cortex. J. Neurosci. 17, 7926–7940.

77. Bellafard, A., Namvar, G., Kao, J.C., Vaziri, A., and Golshani, P. (2024). Volatile working memory representations crystallize with practice. Nature 629, 1109–1117.

78. Trepka, E., Spitmaan, M., Qi, X.-L., Constantinidis, C., and Soltani, A. (2024). Training-Dependent Gradients of Timescales of Neural Dynamics in the Primate Prefrontal Cortex and Their Contributions to Working Memory. J. Neurosci. 44, e2442212023.

79. Fuster, J.M., and Alexander, G.E. (1971). Neuron activity related to short-term memory. Science 173, 652–654.

80. Chen, S., Liu, Y., Wang, Z.A., Colonell, J., Liu, L.D., Hou, H., Tien, N.-W., Wang, T., Harris, T., Druckmann, S., et al. (2024). Brain-wide neural activity underlying memory-guided movement. Cell 187, 676–691.e16.

81. Masse, N.Y., Yang, G.R., Song, H.F., Wang, X.-J., and Freedman, D.J. (2019). Circuit mechanisms for the maintenance and manipulation of information in working memory. Nat. Neurosci. 22, 1159–1167.

82. Dana, H., Chen, T.-W., Hu, A., Shields, B.C., Guo, C., Looger, L.L., Kim, D.S., and Svoboda, K. (2014). Thy1-GCaMP6 transgenic mice for neuronal population imaging in vivo. PLoS One 9, e108697.

83. Pachitariu, M., Stringer, C., Dipoppa, M., Schröder, S., Federico Rossi, L., Dalgleish, H., Carandini, M., and Harris, K.D. (2017). Suite2p: beyond 10,000 neurons with standard two-photon microscopy. bioRxiv, 10.1101/061507. doi: https://doi.org/10.1101/061507.

84. Benjamini, Y., and Hochberg, Y. (1995). Controlling the false discovery rate: a practical and powerful approach to multiple testing. J. R. Stat. Soc. Series B Stat. Methodol. Vol. 57, No. 1, 289–300.

